# Caprin-1 binding to the critical stress granule protein G3BP1 is regulated by pH

**DOI:** 10.1101/2021.02.05.429362

**Authors:** Tim Schulte, Marc D. Panas, Lucy Williams, Nancy Kedersha, Jonas Simon Fleck, Timothy J.C. Tan, Anders Olsson, Ainhoa Moliner Morro, Leo Hanke, Johan Nilvebrant, Kim Anh Giang, Per-Åke Nygren, Paul Anderson, Adnane Achour, Gerald M. McInerney

**Author notes:** These authors contributed equally (Tim Schulte, Marc D. Panas, Adnane Achour, Gerald M. McInerney).

## Abstract

G3BP is the central hub within the protein-RNA interaction network of stress-induced bio-molecular condensates known as stress granules (SG). The SG-associated proteins Caprin-1 and USP10 exhibit mutually exclusive binding to the structured NTF2-domain of G3BP1, thereby regulating G3BP1-mediated condensation, but with opposite effects: Caprin-1 promotes but USP10 inhibits SG formation. Herein, we present the crystal structure of G3BP1-NTF2 in complex with a Caprin-1 derived short linear motif (SLiM), which provides a molecular understanding for the mutually exclusive binding of USP10 and Caprin-1 to G3BP1. Caprin-1 but not USP10 contacts two G3BP1-NTF2 histidine residues, which was confirmed using biochemical, biophysical and cellular biological binding assays. G3BP1/Caprin-1 interactions disrupted via point mutations resulted in fewer and smaller SG condensates. In addition, biochemical binding assays demonstrated reduced binding of Caprin-1 to G3BP1 at lower pH values. Finally, ratiometric pH sensitive measurements of SGs revealed a substantial drop in pH compared to the adjacent cytosol, suggesting that reduced pH can fine-tune and regulate the G3BP1-mediated interaction network via a NTF2-mediated pH-sensitive SLiM-selection mechanism.

## Introduction

Stress granules (SGs) are micron-sized membraneless compartments in eukaryotic cells that are dynamically induced upon environmental and biotic stresses such as oxidation and viral infections ^1–3^. Dysregulated SGs are found in various diseases including cancer, viral infections and neurodegeneration ^4–8^. SGs and other similar membraneless organelles are commonly referred to as bio-molecular condensates (BMCs) ^9^. BMCs result from liquid-liquid phase separation, a process that is governed by heterotypic multivalent interactions between multi-domain proteins and nucleic acids ^10^. The composition and concentrations of SG components are evolutionarily shaped to separate into a condensed phase upon stress and their partitioning follows basic thermodynamic principles ^10–12^. SGs contain mainly non-translating mRNAs and RNA-binding proteins as well as additional proteins affecting their functions ^3,13–18^. The biomolecular composition and interaction networks of SGs are distinct from other BMCs, and also differ based on the type of applied stress or investigated cell type^13,15,19^

Three recent publications have elegantly demonstrated the key role of G3BP1 and G3BP2 (both jointly referred to as G3BP) as effector molecules that mediate SG formation ^20–22^. The authors combined systems biology, *in vitro* reconstitution systems, biochemical assays, cell biology and network theory to identify G3BP as the central hub and regulator of SG formation in selected eukaryotic cell lines. These studies not only confirmed numerous previous studies about the important role of G3BP in SG formation but also suggested a conceptual framework for G3BP-mediated condensation that is based on network theory ^4, 21, 23–27^. According to this theory, macromolecules are defined as node, bridge, cap or bystander molecules depending on the number of interaction sites, *i.e.* equal to or more than three, two, one and zero, respectively. G3BP acts as central node that possesses all the features required to drive phase separation and biomolecular condensation: homotypic oligomerization and heterotypic interactions mediated through the structural NTF2-like domain (hereafter referred to as NTF2), nucleic acid binding through the RRM and RGG domains as well as fine-tuning of SG formation through charged low complexity or intrinsically disordered regions (LCR/IDR) ^16,20–22,28,29^. Caprin-1 and USP10 were identified as prominent regulators of G3BP-mediated condensation that promote and prevent SG formation ^9,20,23–27^. Their disparate roles are explained in the described network theory which posits that USP10 acts as valence cap because it lacks both RNA-binding and oligomerization domains, but efficiently limits NTF2-mediated network interactions via its short linear FGDF motif ^21^. Conversely, Caprin-1 acts as a bridge due to oligomerization and RNA-binding domains as well as its interaction with NTF2, via a short linear motif (SLiM) that is distinct from the biochemically and structurally well-defined FGDF motif^21,28,30,31^.

G3BP-mediated SG condensation is triggered and regulated by electrostatic changes, but also other post-translational modifications such as *e.g.* phosphorylation ^20–23,32–35^. In G3BP1, the phosphorylation status of the two serine residues Ser-149 and Ser-232 within the acidic region may tune the threshold concentration of G3BP1 that is required for condensation ^20–22^. A recent biophysical and functional *in vitro* deadenylation and translation study established that Caprin-1 and fragile X mental retardation protein (FMRP)-derived intrinsically disordered regions (IDRs) only phase-separated upon phosphorylation of either partner, and that the resulting phosphoprotein controlled deadenylation and translation activities due to differences in the nano-scale organization of the resulting condensates ^32^. While phosphorylation adds a negative charge to the modified protein, changes in pH rapidly and reversibly alter the electrostatic surfaces of interacting molecules, exemplifying a simple and evolutionarily conserved post-translational modification ^36^. As G3BP1 adopts a closed conformation at neutral pH, it is open at acidic pH due to electrostatic repulsion between the acidic and basic RNA-binding RGG regions, thus decreasing its saturation concentration for condensation ^22^.

The pivotal role of NTF2 as an interaction hub is exemplified by studies demonstrating that Old World alphaviruses target NTF2 in order to recruit G3BP and associated 40S ribosomal subunits to the vicinity of viral cytopathic vacuoles (CPVs). For this purpose, Semliki Forest virus (SFV) and chikungunya virus express the non-structural protein 3 (nsP3) comprising a duplicated FGDF motif to outcompete USP10 for NTF2-binding ^25,28,31,37^. The multi-domain protein nsP3 contains two FGDF motifs within a largely disordered hypervariable domain (HVD) for multivalent protein interactions, and two structured domains with possibly enzymatic and eventual RNA-binding activities ^31,38^. Strikingly, recruitment of G3BP to CPVs via nsP3 results in the appearance of BMCs that are reminiscent of SGs but have translational activity ^37^.

In this work, we present the crystal structure of the central hub G3BP-NTF2 in complex with a Caprin-1-derived SLiM, providing a molecular understanding of the mutually exclusive binding of USP10 and Caprin-1 to NTF2. The structure revealed an unanticipated ligand selection mechanism mediated by histidine residues on the surface of NTF2. Biochemical and biophysical binding assays, in which these histidine residues were substituted to a selection of amino acids, demonstrated that USP10 and Caprin-1 are finely tuned competitors for NTF2 binding. Importantly, stress-induced BMCs were reduced in number and size in eukaryotic cells expressing NTF2- and Caprin-1 mutants Furthermore, we demonstrate that the Caprin-1/G3BP1 interaction is pH sensitive. Finally, we showed that SGs are themselves of lower pH than adjacent cytosol, suggesting a mechanism for regulation of G3BP ligand selection and fine-tuning of SGs.

## Results

### The YNFI motif of Caprin-1 and the FGDF motif of nsP3/USP10 bind within the same hydrophobic pocket in G3BP1-NTF2

Crystals that were obtained by co-crystallizing G3BP1-NTF2 with a synthesized peptide corresponding to residues 356 – 386 of Caprin-1 (Caprin-1^356-386^) diffracted to a maximum resolution of about 1.9 Å, with diffraction spots of I/σ > 2 extending to about 2.1 Å (Figure S1, Table S1). Six G3BP1-NTF2 molecules were identified in the asymmetric unit arranged as NTF2-dimers (Figure S2). In agreement with previous studies ^31,39^, each NTF2 domain is composed of three α-helices αI– αIII all lined up against a β-sheet created by the five β-strands βI–βV (Figure 1A). The overall conformation of G3BP1-NTF2 is very similar to the previously determined crystal structure of the G3BP1-NTF2/SFV-nsP3^449–473^ complex ^31^ with minimal root-mean square deviations in the 0.3-0.6 Å range. Larger structural deviations were only observed in loop regions. The initial discovery map revealed clear density for 20 of the 31 residues of the Caprin-1^356-386^ peptide that bound to one of the three NTF2 dimers (Figures 1B). The final model was refined to R and R_free_ values of 20.8% and 25.2%, respectively (Figures 1B and S2).

**Figure 1.**
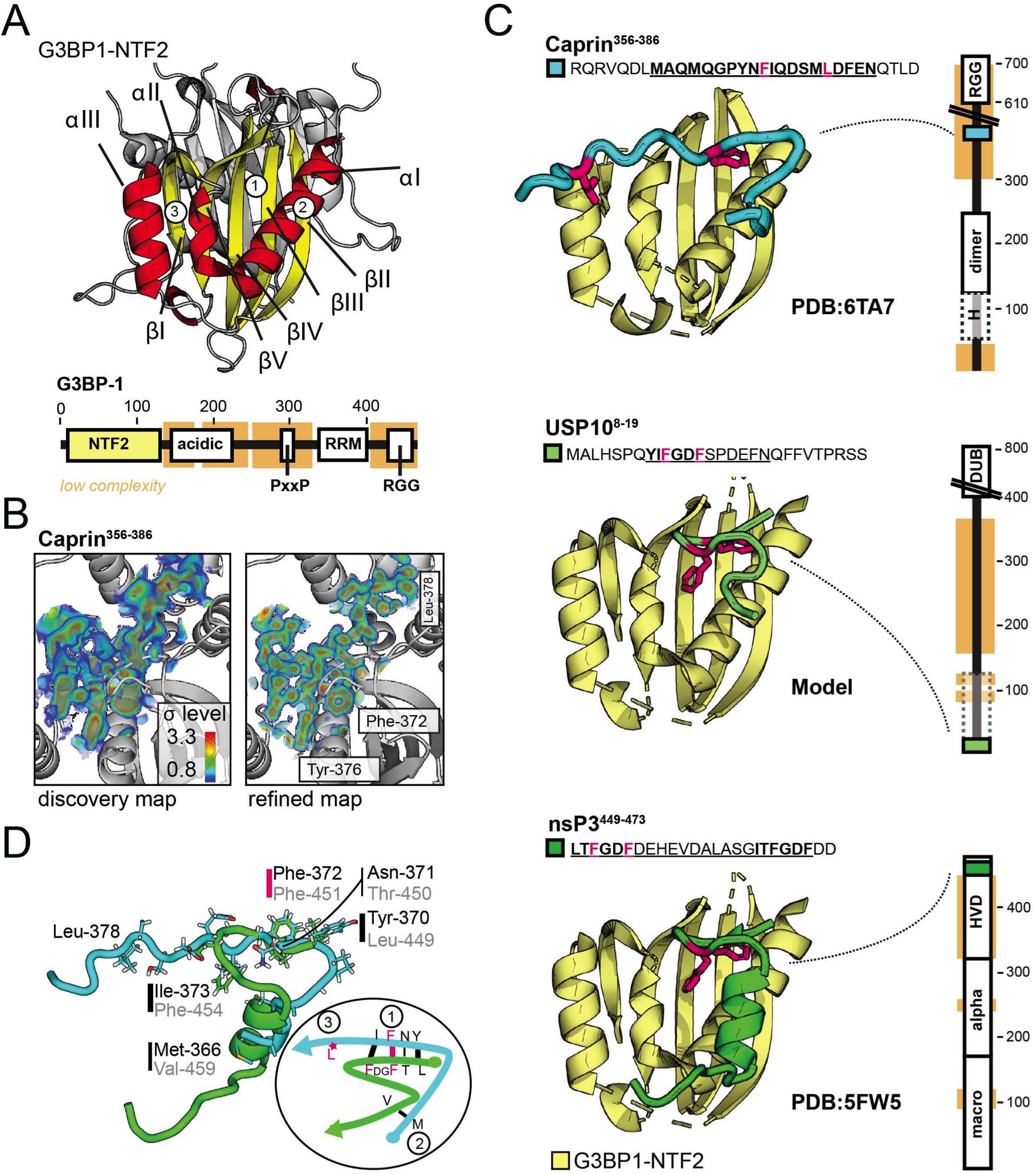
Structural basis for mutually exclusive binding of USP10 and Caprin-1 to G3BP1-NTF2. (A) The structure of the NTF2 domain comprises three α-helices αI–αIII and a β-sheet created by five β-strands βI–βV. Helices and strands are in yellow and red, respectively. The second subunit of the NTF2 homodimer is shown in grey, behind. Full-length G3BP1 (Uniprot: Q13283, scheme below) contains globular NTF2 and RRM domains, as well as structurally disordered regions with low amino acid complexity, including a Glu-rich acidic region, an arginine–glycine–glycine (RGG) box RNA-binding motif and a Proline-rich SH3-binding motif (PxxP) ^29^. The NTF2-like domain is in yellow and residue numbers are indicated above. The three binding sites contacted by Caprin-1^356- 386^ are indicated. (B) The initial 2mFo-DFc “discovery” map revealed 20 out of the 31 residues of the Caprin-1^356-386^ fragment. After model building the map features were clearly improved. G3BP1-NTF2 is shown in grey, density maps are visualized as volumes on depicted color-ramps. See also Figures S1 and S2 as well as Table S1 for crystallographic quality indicators. (C) For simplicity only helix and strand elements of NTF2 are drawn in yellow. Structural models of Caprin-1^356-386^ (PDB: 6TA7, crystal structure), USP10^8-19^ (*in silico* homology model) and nsP3^449–473^ (PDB: 5FW5, crystal structure) are shown in cyan, and green, respectively. See also Figure S3 for USP10^8-19^ model generation. The two phenylalanine residues of LTFGDF as well as Phe-372 and Leu-378 of Caprin-1^356-386^ are highlighted in pink since these were substituted to alanine residues in G3BP1 non-binding mutants for biophysical measurements. Domain organizations illustrate that the NTF2-binding SLiMs are derived from disordered protein segments. Human Caprin-1 (Uniprot ID: Q14444) comprises N-terminal helical regions and a dimerization domain, as well as a C-terminal RNA-binding RGG domain ^24, 30,32^. Human USP10 (Uniprot Q14694) comprises a C-terminal deubiquitinase domain (DUB), long low complexity segments, and a N-terminal domain (dotted lines) that interacts with p53 but also comprises a PAM2 motif at position 78-95 ^28,82,83^. The FGDF-comprising SLiM is at the N-terminus and highlighted in green. SFV nsP3 (Uniprot P08411) is composed of two conserved macro- and Zn-binding domains and a non-conserved hypervariable domain (HVD) comprising two FGDF motifs shown in green ^31,38^. The low complexity NTF2-binding segment is highlighted in cyan. (D) Structures of Caprin-1^356-386^ and nsP3^449–473^ are superimposed in cyan and green, respectively. The scheme highlights the residues of the shared binding regions of Caprin-1^356-386^ and nsP3^449–473^. The third binding region comprising Leu-378 is unique to Caprin-1^356-386^.

A structural comparison of the NTF2/Caprin-1^356-386^ and NTF2/nsP3^449-473^ crystal structures revealed that the YNFI(Q) segment of Caprin-1^356-386^ binds to the same hydrophobic G3BP1-NTF2-pocket (site 1) that was previously identified for the (LT)FGDF motif of nsP3^449-473^ (Figure 1C). Strikingly, the phenylalanine residue Phe-372 in Caprin-1^356-386^ adopts a conformation almost identical to that of residue Phe-451 in nsP3^449-473^ (Figure 1C, compare top, bottom). Although Caprin-1^356-386^ and nsP3^449-473^ cover the same secondary NTF2 binding site (site 2), their N- and C-termini are aligned in opposite directions. More importantly, Caprin-1^356-386^ binding extends to a third NTF2-binding site (site 3) that is not covered by nsP3 (Figures 1C and 1D). Due to its high sequence homology with the (LT)FGDF motif of nsP3^449-473^, the (YI)FGDF-motif of USP10^8-19^ is also most probably positioned at sites 1 and 2, but not site 3, as visualized in a homology-based structural model (Figures 1C and S3).

### The entire length of Caprin-1^356-386^ binds G3BP1-NTF2 with low μM affinity

In order to test the binding predicted by the crystal structure, we performed alanine mutagenesis on residues Q360 to E381 in a full-length Caprin-1 expression construct. We generated U2OS cells lacking endogenous Caprin-1 using CRISPR/Cas9 technology (Figure S4). Caprin-1 mutant constructs were transiently expressed in Caprin-1 KO cells, then immunoprecipitated to determine binding to G3BP1 (Figure 2A). The results demonstrated that residues all along the Caprin-1 fragment were required for efficient binding, in particular site 1 contact-residues Tyr-370, Phe-372 and Ile-373, site 3 contact-residues Ser-376, Met-377, Leu-378, as well as site 1/site 3-contact-residue Gln-374. In addition, the site 1 associated but non-contact residue Gly-368 was also critical for the interaction, likely conferring flexibility to the peptide backbone. It should be noted that substitution of Leu-362, Met-363, Asp-379 and Glu-381 also abolished or reduced binding, although these residues were not resolved or identified as contacting residues in the crystal structure. We speculate that their substitution alters the overall conformation of Caprin-1^356-386^. Ala-substitution of site 2 contact-residues Gln-365 and Met-366, site 1 contact-residues Pro-369 and Asn-371, site 3 contact-residues Asp-375 as well as of residues Gln-360, Asp-361 and Phe-380 did not affect binding and were indistinguishable from wild-type.

**Figure 2.**
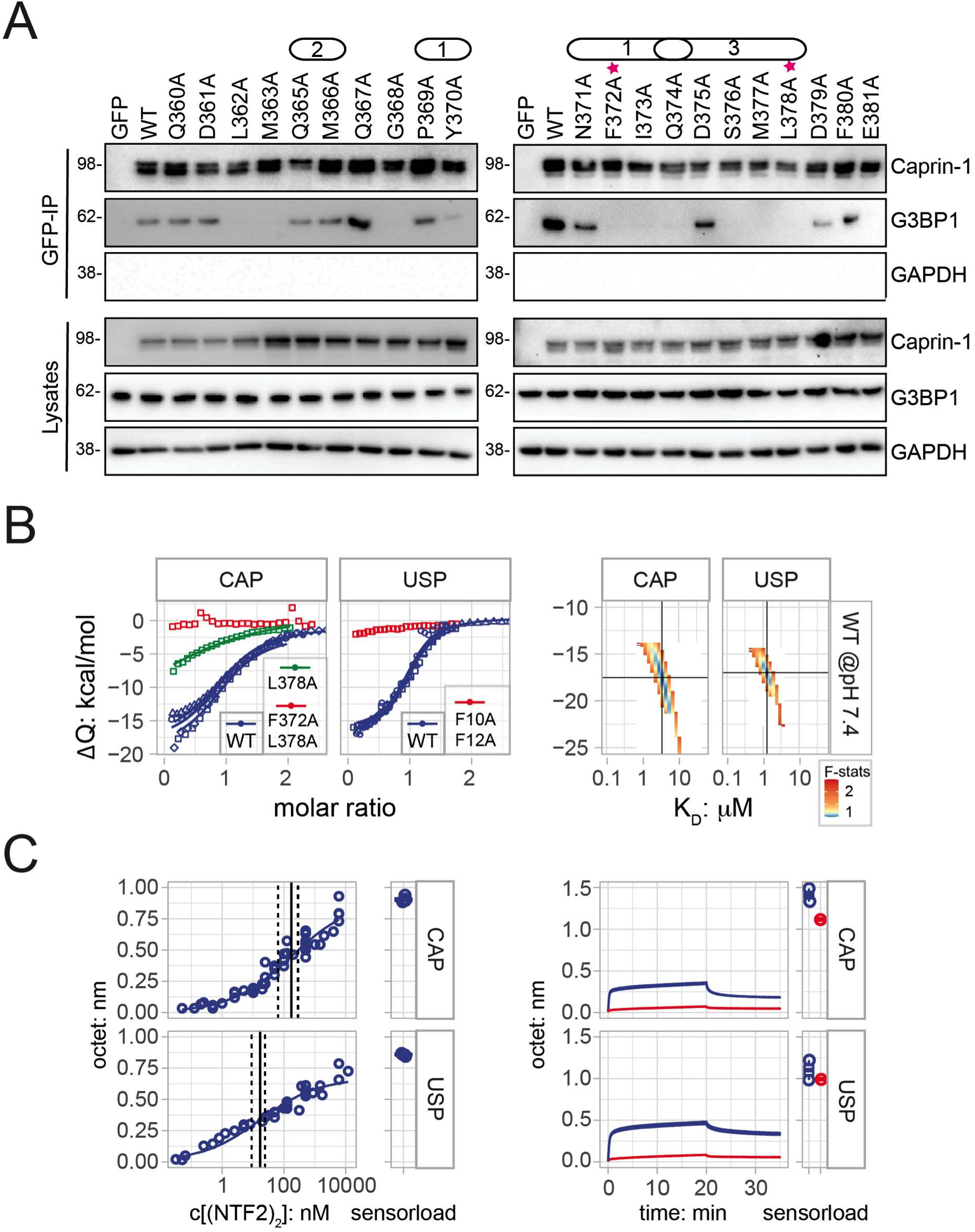
The entire length of Caprin-1^356-386^ binds G3BP1-NTF2 with low μM affinity. (A) GFP-IPs (pulldown) of lysates obtained from ΔCaprin-1 U2OS cells transiently expressing GFP-Caprin-1 WT and mutant constructs demonstrated that almost the entire Caprin-1^356-386^- fragment was required for binding. G3BP1, Caprin-1 and GAPDH were detected by Western blot, cell lysates were applied as protein level controls. Pink stars indicate mutants used for biophysical measurements. Data are representative of at least four repeated experiments. Characterization of Caprin-1 KO cells is described in Figure S4. Identified NTF2 binding sites 1, 2 and 3 are indicated above mutated positions. (B) The integrated heats (data points) and globally fit binding isotherms (lines) from the ITC thermograms (shown in Figure S5) are plotted in panels for the two separate ligands. The twodimensional error surfaces were obtained by F statistics as implemented in Sedphat ^77^ (approximate 1σ-level at a chi-square ratio of 1.1 in yellow). The GFP-Caprin-1^356-386^ to G3BP1-NTF2 binding isotherms (blue) yielded KD and ΔH values of 4 (2 / 6) μM and −18 (−21 / −15) kcal/mol, respectively. The USP10 binding isotherms yielded KD and ΔH values of 1.5 (1/2) μM and −17 (−18 / −16) kcal/mol, respectively. The double mutants (red) yielded minor integrated heats at baseline while the single-mutant Caprin-1^356-386:L378A^ (green) bound with an estimated affinity of about 20 μM. (C) *left panel:* Dose-response curves were obtained from the kinetic BLI measurements shown in Figure S6. Responses were obtained as pseudo-equilibrium binding levels from the end of each association phase of the NTF2 concentration doses. The resulting dose-response curve was fit to yield apparent EC_50_-values of 20 ± 10 nM and 180 ± 110 nM with h-values of 0.5 ± 0.1 and 0.4 ± 0.1 as well as Octet_max_-values of 0.7 ± 0.1 and 0.9 ± 0.1 for GFP-USP10^1-28^ and GFP-Caprin^356-386^, respectively. CAP and USP data were combined from in total 32 and 40 binding curves, respectively. Sensor-immobilization levels are shown next to the binding curves. *right panel:* In a separate comparative experimental setup, kinetic binding curves demonstrated significantly lower NTF22-binding to GFP-Caprin^356-386:F372A/L378A^ and GFP-USP10^1-28:F10A/F12A^ (red traces) compared to wild-type (blue). Wild-type and mutant sensor surfaces were measured as doublets and singlets, respectively, for each ligand. One remaining sensor surface was subtracted as “blank”. See Figure S6 for control measurements at higher NTF2 concentrations.

Previous studies revealed that Caprin-1 and USP10 compete for G3BP1-binding, suggesting proximal binding sites and similar binding affinities ^2^. Indeed, isothermal titration calorimetry (ITC) experiments revealed similar low micromolar affinities and thermodynamic signatures for both GFP-Caprin-1^356-386^ and GFP-USP10^1-28^ (Figure 2B). Injection of GFP-Caprin-1^356-386^ to NTF2 yielded exothermic heat changes with initial peaks in the 0.5 μcal/s range that returned to baseline during the final injections (Figure S5). The global heterogeneous 1:1 model fits to the derived binding isotherms from four GFP-Caprin-1^356-386^-NTF2 titrations yielded affinity (K_D_) and enthalpy (ΔH) values of 4 ± 2 μM and −18 ± 3 kcal/mol, respectively (Figure 2B). Low micromolar affinity and similar exothermic enthalpy values were also obtained for binding of GFP-USP10^1-28^ to NTF2 from three binding isotherms with K_D_ and ΔH-values of 1.5 ± 0.5 μM and −17 ± 1 kcal/mol, respectively. Based on the co-IP assays and our previous studies ^28,31^, we selected GFP-Caprin-1^F372A/L378A^ and GFP-USP10^F10A/F12A^ as negative control mutants for ITC. Indeed, injection of these double mutants to NTF2 yielded minor heat fluctuations at baseline level (Figures 2B and S5). Titration of single-mutated variant Caprin-1^356-386:L378A^ caused heat changes that were due to an interaction with a significantly reduced affinity in the 20 μM range (Figure 2B).

For complementary kinetic Bio-Layer Interferometry (BLI) measurements, we immobilized the GFP-USP10/Caprin-1 ligands via chemical amino-coupling most likely via the available GFP-lysine residues. This strategy allowed us to apply simple regeneration strategies that should not affect the disordered SLiMs. However, we noticed that G3BP1-NTF2 bound significantly to the negatively charged dextran sensor surface, so this was considered by subtracting NTF2-to-blank binding at each analyte concentration. Thereby we determined apparent EC50-values for binding of NTF2 to GFP-USP10^1-28^ and GFP-Caprin^356-386^ with 20 nM and 180 nM, respectively (Figures 2C and S6A). Obviously, the apparent affinities indicated much stronger binding compared to the ITC-derived values. This overestimation was likely due to dominating electrostatic attraction between the basic G3BP-NTF2 (theoretical pI of about 9) and the acidic sensor surface. Furthermore, the bivalent nature of NTF2 also led to expected avidity and rebinding effects. Importantly, NTF2 bound significantly less to the mutants GFP-USP10^F10A/F12A^/GFP-Caprin-1^F372A/L378A^ at the two tested NTF2 concentrations of 50 and 500 nM (Figures 2C and S6B right panels). Due to the high background binding, fast association kinetics and complicating avidity effects, we did not interpret the binding kinetics.

In conclusion, GFP-Caprin-1^356-386^ and GFP-USP10^1-28^ both bind G3BP1-NTF2 in a similar micromolar affinity range as determined by ITC, with slightly stronger binding by GFP-USP10^1-28^. Importantly, BLI measurements confirmed stronger binding of the USP10-derived SLiM.

### Caprin-1 Leu-378 contacts histidine residues His-31 and His-62 within a third NTF2 binding site outside those covered by USP10 or nsP3^449-473^

A simple residue-level interaction counter highlighted Caprin-1^356-386^-YNFI(Q) and nsP3^449-473^- LTFGDF as the major interaction hot spots (Figures 3 and S7). Since both segments cover essentially the same NTF2-binding sites site 1 and site 2 (Figures 1 and 3), we refer the reader to the supplement for a more detailed description. We only compare the available crystal structure models in molecular detail, but the same major conclusions apply to the highly homologous (YI)FGDF-motif of USP10^8-19^ (Figures 1C and S3). Herein, we focus on site 3 located between helices αII and αIII that uniquely interacts with a stretch of five Caprin-1^356-386^ residues comprising Gln-374 to Leu-378 (Figures 3 and 4A). Residue Gln-374 is the connecting link between site 1 and site 3 and contacts NTF2-residues Phe-124, Tyr-125, Lys-123 and Arg-32. These interactions also comprise hydrogen bonds between the backbone amide groups and the backbone and side-chain guanidine groups of Arg-32, as well as another hydrogen bond between the side-chain amide and the amino-group of Lys-123 (Figure 3). The backbone carbonyl-oxygen of Asp-375 is fixed by a hydrogen bond to the side-chain amide-nitrogen of Gln-58 (hydrogen bond not highlighted, but residues are shown in Figures 3 and 4A). Interestingly, the side-chain of Caprin-1^356-386^ residue Leu-378 is positioned centrally above the triangle that is created by the Cα atoms of NTF2 residues Gln-58, His-31 and His-62 (Figure 3). While the two pH-sensitive imidazole groups have *in silico* estimated pKa-values of 6.1/6.2 in the unbound state, Caprin-1^356-386^ binding shifts the pKa of His-31 to a value of 4.5 due to hydrogen bond formation with Ser-376, while the pKa of His-62 is slightly increased to a value of 6.5 (Figure 4A).

**Figure 3.**
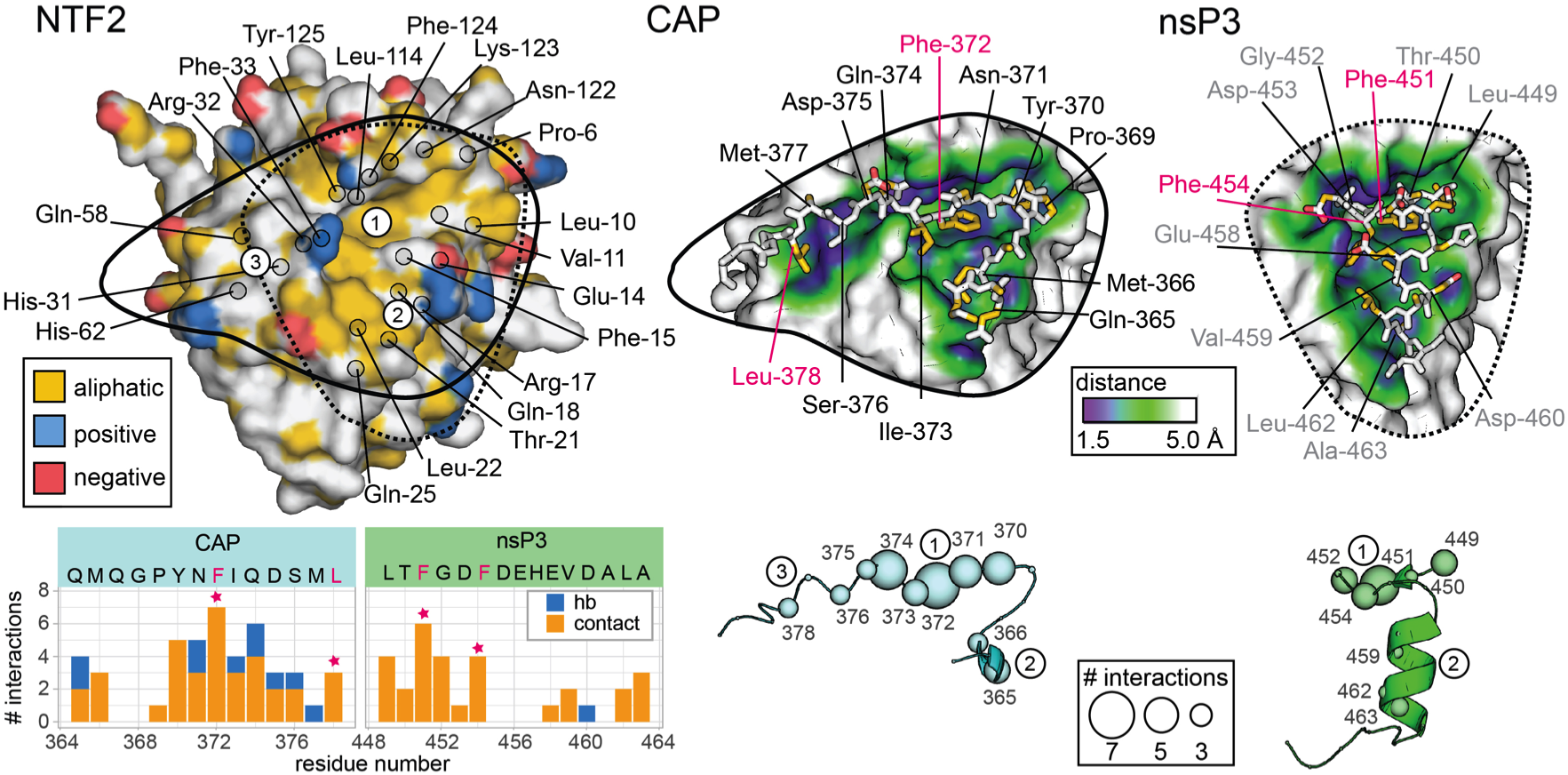
YNFI(Q)- and FGDF cover the same NTF2 binding site, but only Caprin-1^356-386^ extends to a third site between helices α2 and α3. The structural interface overview illustrates that YNFI(Q) and LTFGDF are coordinated by essentially the same NTF2-residues. *top left:* Functional aliphatic, negative and positive groups are colored yellow, red and blue on the NTF2 surface (yrb-colored). Backbone-C_α_ atoms of residues in contact with any of the two ligands are highlighted in black circles. NTF2 binding sites are indicated. *top right:* Caprin-1^356-386^ and nsP3^449–473^ are visualized as yrb-colored sticks on surface patches of NTF2 that are colored according to the depicted distance ramp. *bottom left:* A simple residue-level interaction count highlights the major interaction sites. Contacts and hydrogen bonds identified by Molprobity were summarized and visualized on residue-level as bar-charts colored in orange and blue, respectively. See also Figure S7 for a similar plot based on interaction interface areas. *bottom right:* The total number of contacts of each residue of nsP3^449–473^ and Caprin-1^356-386^ in green and cyan, respectively, is visualized on the depicted C_α_ size scale.

**Figure 4.**
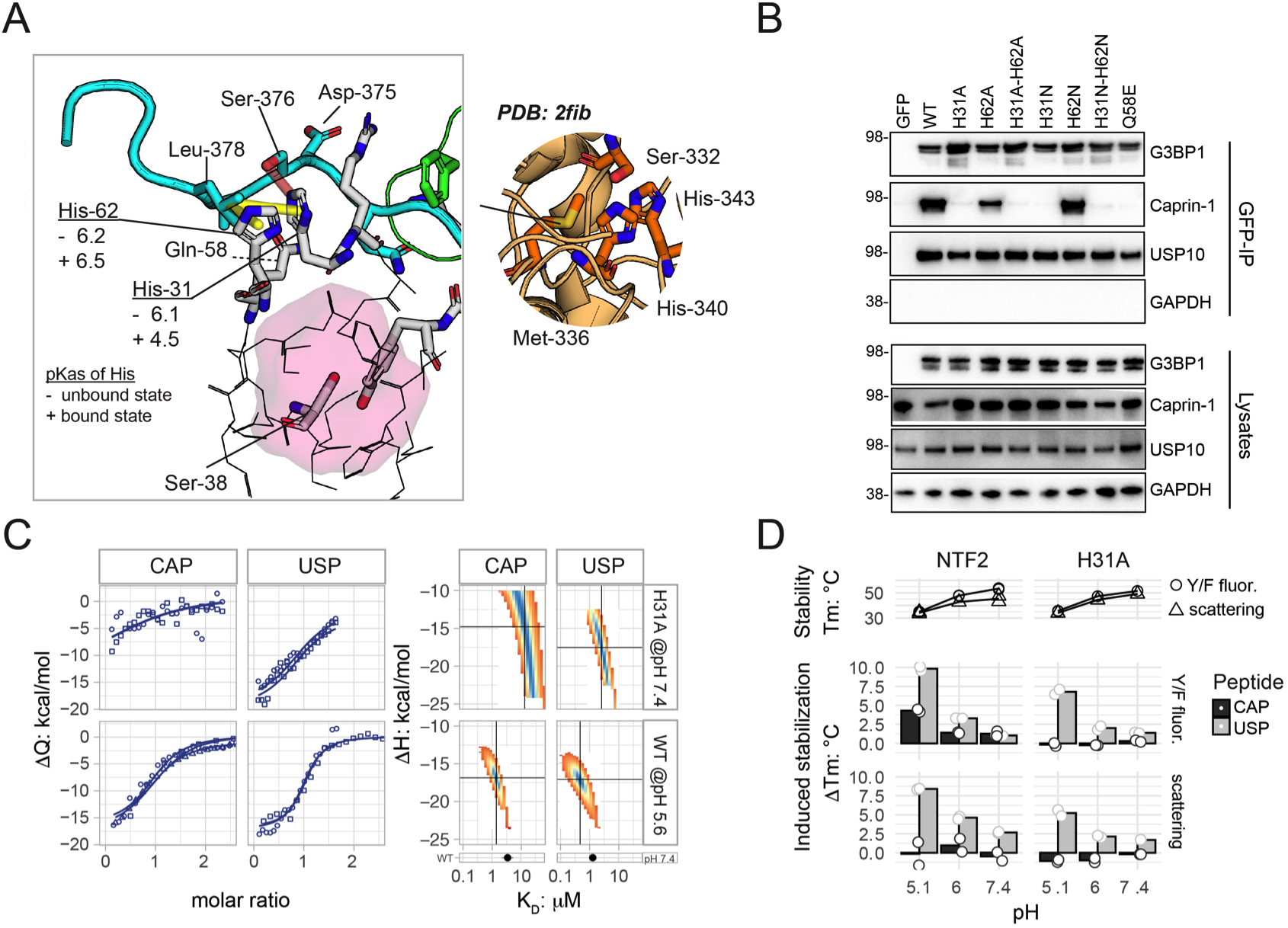
NTF2-His31 contacts Caprin-1 but not USP10. (A) Leu-378 is coordinated by His-31 and His-62 with predicted pK_a_-values of 6.1 and 6.2 in the unbound state (-), respectively. The pKa value of His-31 is predicted to shift to a value of about 4.5 in the Caprin-1 bound state (+) due to the hydrogen bond with Ser-376 and an increased buried surface area. Superimposed Caprin-1 and USP10 ligands are shown in cyan and green, respectively. Important side-chains are highlighted as sticks. Ser-38 and interface residues exclusive to Caprin-1 binding are shown as grey sticks. Residues comprising atoms that lie within a radius of below 8 and 4 Å from the Ser-38 side-chain are shown as thin lines and highlighted by the semi-transparent red surface, respectively. Selected hydrophobic and hydrogen bond interactions are shown as yellow and red semi-transparent dashes, respectively. A similar threedimensional arrangement was identified in the pH-sensitive polymerization site of fibrinogen comprising His-340, His-343, Met-336 and Ser-332 ^46^ (right). (B) U2OS ΔΔG3BP1/2 cells were stably transfected with indicated GFP-G3BP1 mutant constructs. GFP immunoprecipitates and full-cell lysates were analyzed by Western blot for G3BP1, Caprin-1, USP10 or GAPDH. Data are representative of at least five repeated experiments. (C) ITC data were visualized as in Figure 2B. The isotherm of each GFP-USP10^1-28^ and GFP-Caprin-1^356-386^ binding to NTF2-H31A at pH 7.4 (top panel) yielded K_D_-values of 3 μM (1 / 5) and 15 μM (5 / 30) as well as ΔH-values of −18 (−23 / −13) and −16 (−24 / −10) kcal/mol, respectively. The isotherms of wild-type GFP-USP10^1-28^ and GFP-Caprin-1^356-386^ to NTF2 binding at pH 5.6 (bottom) yielded K_D_-values of 0.5 μM (0.3 / 0.7) and 1.4 μM (1.1 / 1.7) as well as ΔH-values of −17 (−19 / −16) and −17 (−18 / −16) kcal/mol, respectively. See also Figure S5 for thermograms. (D) nanoDSF measurements revealed reduced G3BP1-NTF2 stability at acidic pH. This destabilization is compensated better by binding of the USP10-derived SLiM compared to Caprin-1. G3BP1-NTF2 melting temperatures (T_m_) were detected as changes in Tyr/Phe fluorescence (T_m_- F) and scattering (Tm-Sc). ΔT_m_-values were calculated as T_m_(ligand-bound)-T_m_(unbound). Raw data traces and peptide titrations are shown in Figure S8.

Co-IP assays confirmed the importance of these residues for binding of Caprin-1^356-386^ to site 3 of NTF2, since single alanine substitutions of residues Gln-374, Ser-376, Met-377 and Leu-378 abolished binding (Figure 2A). It should be noted that the GFP-Caprin-1^356-386^ L378A mutant bound to NTF2 in our ITC measurements, albeit with a significantly reduced affinity of about 20 μM (Figure 2B). Since histidine residues frequently act as acid-base components within catalytic reaction centers or as pH-sensitive interaction regulators ^40–44^, we applied a protein database (PDB) search strategy ^45^ to identify three-dimensional arrangements similar to the one created between NTF2 residues His-62 and His-31, and residues Leu-378 and -Ser-376 in the Caprin-1^356-386^ fragment. We located a motif comprising His-340, His-343, Met-336 and Ser-332 within the pH-sensitive polymerization site of fibrinogen ^46^. The authors presenting that structure proposed a mechanism whereby ionization of either or both of these histidine residues determines the pH dependence of fibrin polymerization ^46^. Since the NTF2 residues His-62 and His-31 are contacting residues of Caprin-1 but not USP10, we hypothesized that the interaction of NTF2 with Caprin-1 but not that of USP10 might be affected by changes in local pH.

### The NTF2 residue His-31, localized within the third interaction site, selects for Caprin-1 but not USP10

Before testing the pH dependence of the Caprin-1/G3BP1 interaction, we sought to investigate the importance of the third NTF2-binding site for binding to Caprin-1. We generated GFP-G3BP1 constructs in which residues His-31 and His-62 were substituted to alanine or asparagine, and a construct in which Gln-58 was mutated to glutamic acid. We expressed the G3BP1 mutants in U2OS ΔΔG3BP1/2 cells ^2^ and performed immunoprecipitation experiments to determine levels of binding to Caprin-1 and to USP10. Substitution of His-31 to either alanine or asparagine or of Gln-58 to glutamic acid abolished binding to Caprin-1 while binding to USP10 was unaffected (Figure 4B). Substitution of His-62 to alanine markedly reduced binding, but substitution to asparagine did not. Taken together, these data show that G3BP1 residues His-31 and Gln-58 are critical for the interaction with Caprin-1 while residue His-62 plays a minor role. ITC measurements at neutral pH revealed that Caprin-1 bound to NTF2-H31A with a drastically reduced double-digit μM affinity while USP10 bound to NTF2-H31A with an affinity in the 1-5 μM range, slightly higher than the value obtained for wild-type NTF2 (Figures 4C and S5). However, although we hypothesized that a larger fraction of residue His-31 should be protonated at pH 5.6 with reduced NTF2 binding, ITC measurements performed at this pH did not substantiate our hypothesis (Figures 4C and S5). Instead, both interactions were in the low micromolar range with fitted affinity values of 1.4 ± 0.3 and 0.5 ± 0.2 μM, respectively. ITC measurements at pH 5 could not be performed due to NTF2 protein instability. We also performed nano differential scanning fluorimetry (nanoDSF) analysis of G3BP1-NTF2 to reveal reduced NTF2 stabilities at lower pH, with melting temperatures (Tm) dropping from 53 to 48 and 35°C upon shifting the pH from 7.4 to 6.0 and 5.1, respectively (Figures 4D and S8). Interestingly, the USP10^1-28^ peptide stabilized NTF2 significantly better than Caprin-1^356-386^ at the lower pH-values of 6 and 5.1, as the Tm-values were increased by 3 and 10°C, respectively, compared to only 1.5 and 4 °C for Caprin-1^356-386^. In control measurements, NTF2-H31A was stabilized by USP10^1-28^, but not Caprin-1^356-386^ (Figure 4D). The thermal stability of NTF2 was increased marginally by 1 and 1.3 °C when bound to USP10^1-28^ or Caprin-1^356-386^ at pH 7.4 (Figures 4D and S8).

Thus, although our ITC data did not demonstrate lower binding affinities of Caprin-1 to G3BP1-NTF2 at acidic pH, nanoDSF measurements demonstrate the enhanced capacity of USP10 but not Caprin-1 to stabilize NTF2 at acidic pH compared to Caprin-1.

### Reduced condensates in stressed NTF2-H31A mutant cells

Since Caprin-1 and USP10 bind G3BP in a mutually exclusive manner, with positive and negative effects on SG formation, respectively ^2^, we investigated SG formation in U2OS ΔΔG3BP1/2 cells stably expressing GFP-G3BP1-WT as well as H31A, H62A, or Q58E mutants under sodium arsenite (SA) stress and unstressed (mock) conditions. Qualitative fluorescence microscopy images revealed decreased SG numbers and sizes in both G3BP1-H31A and G3BP1-Q58E, compared to G3BP1-WT (Figure 5A). SG numbers and sizes of the H62A mutant were at an intermediate level. We noticed generally lower Caprin-1 levels in condensates of H31A and Q58E cells. These qualitative estimations were confirmed by our high content imaging analysis pipeline in which about 80-90% of in total 3000 GFP-positive cells for each G3BP1 construct were identified as SG-positive upon SA treatment, compared to a false-positive rate of about 15% for mock-treated G3BP-WT cells (Figure 5B). Despite the false-positive SG rate for non-stressed cells, the pipeline clearly differentiated them from stressed cells. Manual inspection of the images showed that mock-treated cells did not contain visible SGs, as expected.

**Figure 5.**
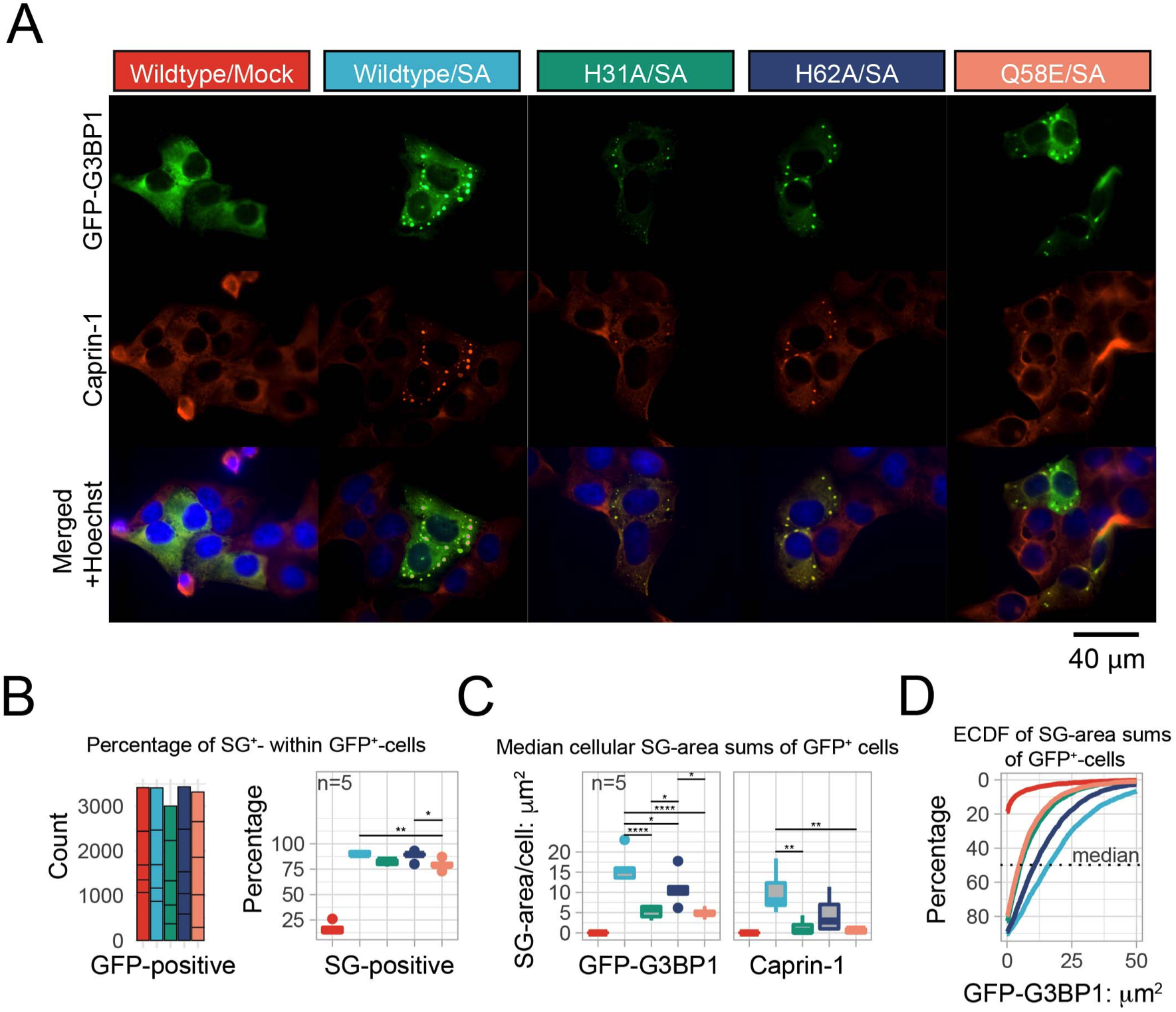
NTF2-His31 mutants form fewer stress granules with reduced Caprin-1 content. (A) Representative microscopy images of ΔΔG3BP1/2 U2OS cells stably expressing indicated G3BP1-wildtype, -H31A, -H62A and -Q58E were stressed with sodium arsenite (200 μM) for 1h and compared to non-stressed (mock) cells. (B-D) Analysis summary of high-content microscopy imaging (see related Figure S9). SG areas in GFP-positive cells were summed separately for the two GFP-G3BP1 and Caprin-1 channels. In our analysis we refer to the cellular SG-area as the summed area of all SGs within a single cell, and not the area of a single SG. (B) Total counts of GFP-positive cells per experiment yielded similar numbers of around 3000 cells for each population. The percentages of GFP-G3BP1 SG-positive cells of each population were determined separately for each experiment (n = 5) and visualized as a boxplot. Significance p-levels were obtained from post-hoc ANOVA Tukey HSD: *_≤ 0.05, ** ≤ 0.01, *** ≤ 0.001, **** ≤ 0.0001. The significance labels for the comparison between stressed and unstressed mock cells were removed. (C) Median values of the cellular SG area sums were extracted from each experiment to estimate their experimental variation in the GFP-G3BP1 and Caprin-1 channels (see also Figure S9C). (D) The GFP-positive cell populations were combined from all experiments and represented as ECDF for a direct comparison in a single plot. The y-axis represents the percentage of remaining cells: about 80-90% of stressed cells, while 15% of non-stressed cells had SG area sums > 0 μm^2^. The median is shown as a dotted line.

To reveal differences at the single-cell level, the areas of all SGs in each cell were summed separately for the two GFP-G3BP1 and Caprin-1 channels (Figure S9). For better comparison between the different mutants, and to combine SG size and number, we quantified the total SG-area per cell (in the following referred to as SG-area). A two-dimensional visualization of this cellular SG-area confirmed our initial qualitative estimate that SA-stressed G3BP1-WT cells had larger total SG-areas per cell compared to H31A or Q58E, while H62A-cells displayed intermediate-sized SG-areas (Figures 5C and S9B). Median values of the cellular SG-areas were derived from each experiment to estimate their experimental variation in the two GFP-G3BP1 and Caprin-1 channels (Figure S9C), yielding mean values of 16.5 ± 4.0 and 10.0 ± 5.5 μm^2^, 5.5 ± 1.5 and 1.5 ± 2.0 μm^2^, 11.0 ± 4.0 and 4.5 ± 4.5 μm^2^, as well as 5.0 ± 1.0 and 0.5 ± 0.5 μm^2^ for G3BP1-WT, -H31A, -H62A as well as -Q58E, respectively (Figure 5C). This analysis demonstrated significantly larger SG areas in the GFP-channel for stressed G3BP1-WT cells compared to all other cells. Increased condensate areas resulted from higher numbers and slightly enlarged areas of individual granules in each cell, e.g. G3BP1-WT had 7.5 ± 1.5 SGs/cell with area/SG of 2.0 ± 1.5 μm^2^, while G3BP1-H31A cells had 4.0 ± 1.0 SGs/cell with area/SG of 1.5 ± 2.0 μm^2^ (Figure S9D). Unstressed mock-treated cells had median SG-areas of zero in both channels (Figure 5D). The G3BP1-H62A-mutant cells had significantly larger SG-areas than both H31A and Q58E cells. The GFP-G3BP1 SG-areas of the combined datasets were also visualized as empirical cumulative distribution functions (ECDF) to compare the cell populations directly in a single plot (Figure 5D), revealing the same trends: WT > H62A > H31A ≈ Q58E > Mock. Similar trends were observed for the Caprin-1 channel, but it should be noted that the determined SG-areas were generally smaller (Figure 5C). The Caprin-1-SG-areas of WT, H31A, H62A and Q58E cells were reduced by 40, 75, 60 and 85%, respectively, compared to the GFP-G3BP1 channel. The capacity of the cells to efficiently recruit Caprin-1 into condensates correlated with condensate size, and closely matched the G3BP1/Caprin-1 binding pattern that was observed in our biochemical G3BP1/Caprin-1 co-IP assays (Figure 4B). In agreement with previous studies ^2,20^, our data thus reveal that Caprin-1 positively affects, but is not essentially required for, G3BP1-mediated condensation, recognized herein as reduced condensate formation in G3BP1-site 3 mutated cells.

### Lowered pH, detected within stress granules, destabilizes Caprin-1/G3BP1 but not USP10/G3BP1 complexes

To investigate whether G3BP1-His-31 and His-62 are involved in pH-dependent regulation of Caprin-1 binding, we adapted our established co-IP assays. Immunoprecipitated GFP-Caprin-1-WT associated complexes were washed with buffers of pH values 5.6, 6.1, 7.0 and 7.4, and proteins remaining in complex were detected by western blot. The stability of GFP-Caprin-1-WT/G3BP1 was ^32,34,47^ significantly reduced at acidic pH, while the interaction of the Caprin-1 binding protein FMR1 ^32,34,47^ was stable and non-significantly enhanced at pH 6.1 (Figures 6A and 6B). As a control, we designed a chimeric Caprin-1-FGDF construct, in which the putatively pH-sensitive Caprin-1^360–383^ NTF2-binding-motif was replaced by the USP10^1-25^ NTF2-binding motif. This chimera associated more strongly with G3BP1 and the formed complexes were not sensitive to acidic pH washes. Binding of FMR1 was not changed by alterations in buffer pH (Figures 6A and 6B). Thus, acidic pH only destabilized the G3BP1-NTF2/Caprin-1^360-383^ complex but not G3BP1-NTF2/Caprin-1-FGDF.

**Figure 6.**
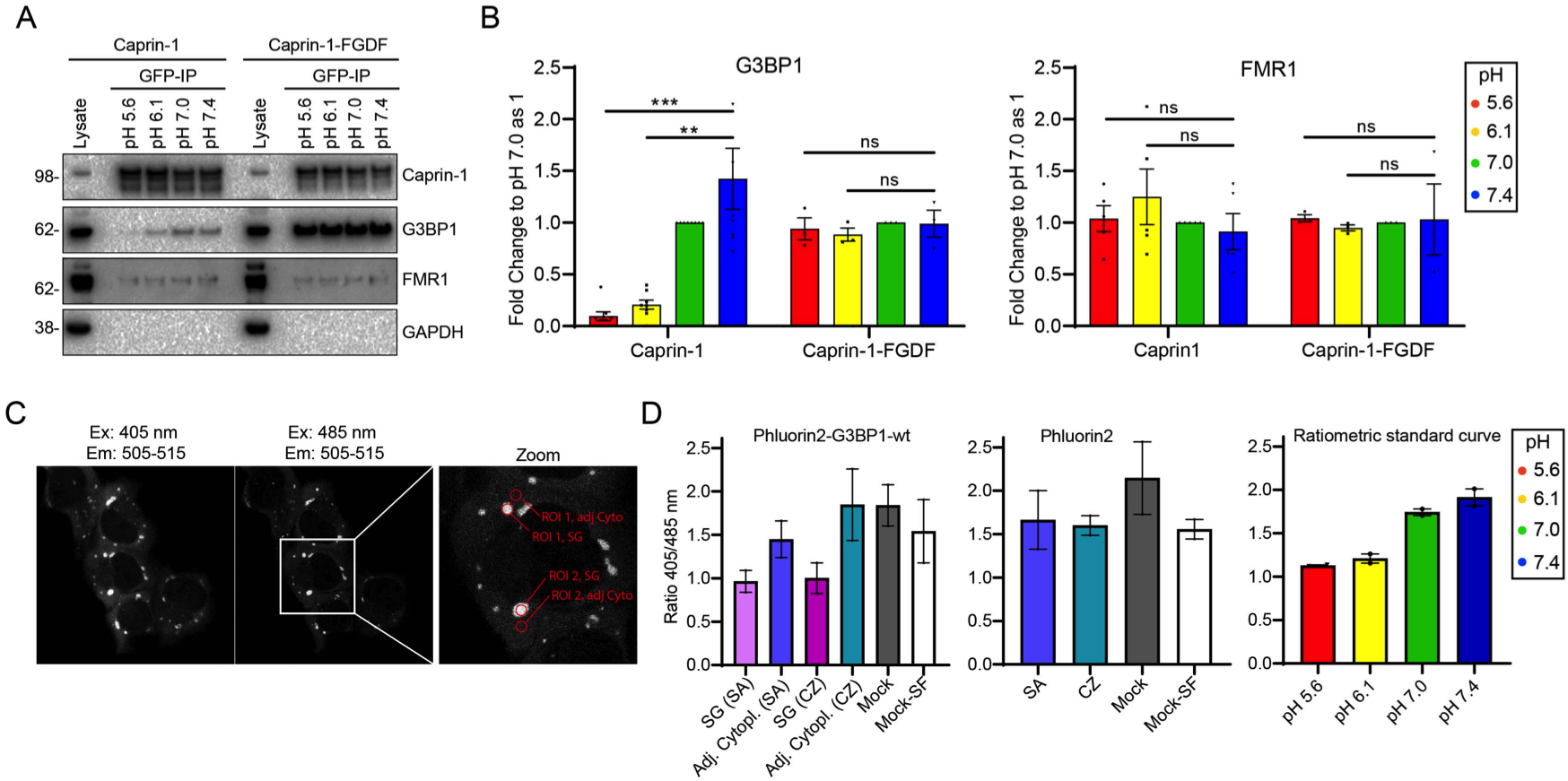
Low pH is observed within SGs and destabilizes Caprin-1/G3BP1 but not Caprin-1-FGDF/G3BP1. (A) U2OS ΔCaprin1 cells were stably transfected with GFP-Caprin-1 WT or GFP-Caprin-1-FGDF. GFP fusions were precipitated using GFP-Trap agarose and washed with pH-adjusted buffers. Immunoprecipitates and full cell lysates were analyzed by Western blot for Caprin-1, G3BP1, FMR1 or GAPDH. Data are representative of at least three repeated experiments. (B) Western blots signals were quantified by densitometry and the ratios of coimmunoprecipitated G3BP1 or FMR1 protein relative to GFP-Caprin-1 WT or GFP-Caprin-1-FGDF were determined. Data shown are mean ± SEM and are analyzed using an unpaired t test. ns: non-significant, **, P < 0.01; n = 8 for GFP-Caprin-1 WT and n=3 for GFP-Caprin-1-FGDF IPs. (C) pH-dependent ratiometric changes of pHluorin2 were monitored to estimate the intracellular pH within SGs and the adjacent cytoplasm. Fields of live stressed ΔΔG3BP1//2 U2OS cells stably expressing pHluorin2-G3BP1-wt were sequentially excited at two excitation wavelengths of 405 and 485 nm, and the emission was detected between 505-515 nm. The enlarged image (zoom) exemplifies our procedure to derive fluorescence intensities from regions of interest (ROI) within SGs and the adjacent cytoplasm. (D) Images of ΔΔG3BP1//2 U2OS cells stably expressing pHluorin2-G3BP1-wt or pHluorin2 were analyzed as described in (C), then 405/485 nm ratios were calculated and plotted. To estimate approximate absolute pH values, we also applied the same procedure on beads with bound pHluorin2 (see Figure S10) and analyzed the images as described in (C), then 405/485 nm ratios were calculated and plotted. Two biological independent experiments were analyzed.

The pH-dependence of the Caprin-1/G3BP1 interaction prompted us to ask whether SGs are acidic. Recent studies propose that pH-dependent conformational changes of the acidic IDR of full-length G3BP1 fine-tune phase-separation and SG formation ^20–22^. While pH has been previously proposed as a general evolutionarily conserved fine-tuning mechanism for yeast SG regulation^11^, we were unaware of any study measuring the pH within mammalian SGs. To assess this, we fused the pH-sensitive GFP variant pHluorin2 ^48^ to G3BP1-WT and used that fusion protein to estimate the pH in SGs relative to equally sized areas of adjacent cytoplasm in live ΔΔG3BP1/2 U2OS cells (Figure 6C). Ratiometric pHluorin2 exhibits a bimodal fluorescence excitation spectrum with peak maxima at 395 and 475 nm when the emission is recorded at the fluorescence peak maximum at 509 nm. Acidic pH decreases the fluorescence excitation peak at 395 nm, but increases that at 475 nm, providing a means to monitor intracellular pH changes. Before we tested pHluorin2 in live cells, we recorded a standard curve of bead-immobilized GFP-pHluorin2 in buffers with distinct pH-values (Figures 6D and S10). We note that the standard curve should not be used to derive absolute pH-values within living cells due to nonidentical surrounding environments and slightly altered image acquisition settings between the bead and cellular measurements. Nevertheless, the standard curve sets a useful reference frame for the expected pHluorin2 signal in our setup. We observed a decrease in the 405/485 ratio from about 1.9 at pH 7.4 to about 1.1 at pH 5.6 for the bead measurements, as expected ^48^ (Figure 6D). Similarly performed analyses for the cytoplasm of stressed and non-stressed cells yielded fluorescence ratios of 1.5 and 1.8, respectively. Both sodium-arsenite- and clotrimazole-induced stress granules (SG/SA and SG/CZ, respectively) had significantly reduced emission ratios with values of about 1, indicating considerably more acidic environments within these condensates relative to the adjacent cytoplasm. These observations suggest that a slightly more acidic environment within SGs could fine-tune binding of Caprin-1 to G3BP.

## Discussion

The crystal structure of the NTF2/Caprin-1^356-386^ complex at 2.1 Å resolution provides a molecular explanation for the mutually exclusive binding of Caprin-1 and USP10 to G3BP1-NTF2 ^2,20–22^. The Caprin-1^356-386^-derived central binding motif YNFI(Q) occupies the same hydrophobic binding pocket on the NTF2 surface as do the USP10- and alphavirus nsP3-derived (YI/LT)FGDF motifs ^31,39^. Both the Caprin-1 and USP10 /nsP3 motifs share a central phenylalanine that serves as the main aromatic anchor residue, adopting the same conformation in both ligands (Figure 1). Both binding motifs are within IDRs and these interactions therefore exemplify motif-based protein-protein interactions that are characterized by binding of SLiMs to folded protein domains ^49,50^. SLiMs are on average 6-7 amino acids in length and contain only 3-4 core positions as main anchor residues. Due to their small interaction interface areas such interactions are typically transient and in the low-affinity 1-500 μM range. Although a crystal structure of the USP10-derived SLiM is not available, structural modeling based on the highly homologous SFV-nsP3^449–473^ derived (LT)FGDF motif suggests almost identical binding interfaces for both nsP3 and USP10 (Figure 1 and S3). The nsP3^449–473^ and Caprin-1^356-386^ motifs cover total interface surface areas with NTF2 of 740 and 860 Å^2^, respectively ^31^. Their central LTFGDF and YNFI(Q) motifs cover areas of 550 and 525 Å^2^ which are close to the 500 Å^2^ value that is typically observed for SLiM interface areas ^49^. Both USP10 and Caprin-1-derived SLiMs bind NTF2 with single digit micromolar affinities as determined by ITC, placing both ligands at the higher affinity end of typical SLiM interactions.

Importantly, the structure revealed that the Caprin-1^356-386^ residue Leu-378 contacts the pH-sensitive G3BP1-NTF2 residues His-31 and His-62, both localized within an extended binding site (site 3) not bound by either nsP3 or USP10. This discriminative contact is fixed by a hydrogen bond between the imidazole-group of the NTF2 residue His-31 and the hydroxyl group of the Caprin-1 residue Ser-376. Notably, a structurally similar interaction was described for the pH-sensitive polymerization site of fibrinogen and proposed as a possible regulatory mechanism ^46^. Our biophysical and biochemical assays demonstrated that alanine-substitutions of NTF2-His-31 abolished binding to Caprin-1^356-386^ but only minimally affected the interaction with USP10. Further support for a ligand selection mechanism was previously provided by Sanders and co-workers who serendipitously discovered that a G3BP1-NTF2-S38F mutant binds USP10 but not Caprin-1 ^21^. G3BP1-NTF2 Ser-38 is localized in the vicinity of site 3 (Figure 4A), and we speculate that its substitution to a phenylalanine causes larger-scale perturbations preventing Caprin-1 binding.

In line with the SG-network theory and recent phase-separation studies ^20–22^, mutations within the third NTF2-binding site led to decreased condensation in stressed cells, as demonstrated by high-content imaging of ΔΔG3BP1/2 cells stably expressing the WT GFP-G3BP1 construct as well as the H62A, H31A and Q58E G3BP1 variants (Figure 5), resulting in the following sequence of SG-area sum per cell: WT > H62A > H31A ≈ Q58E > Mock (null). The condensation sequence followed the same binding pattern that was observed in Caprin-1:G3BP1 IP-binding assays (Figure 4B). Thus, reduced binding of the node G3BP1 with its bridge Caprin-1 reduced condensation significantly. However, we cannot exclude that the introduced mutations also affect other bridge or node molecules, such as UBAP2L. It should also be noted that Caprin-1 is recruited into SGs independently of G3BP1^24,32,47^ due to its interaction with mRNA and FMR1 ^24,32,47^, but this recruitment is significantly reduced in G3BP1-site 3 mutants, suggesting the partitioning of Caprin-1 into SGs is to a large extent driven by its interaction with G3BP. Furthermore, the reduced SG condensation of the different mutants could also be attributed to more effective competition with USP10, which acts as a valency cap and blocks SG formation when overexpressed ^2,20–22^.

Determination of the crystal structure allowed us to identify histidine residues in NTF2 that contact Caprin-1, which prompted us to test the pH-dependence of this interaction, especially since pH-induced conformational changes of G3BP1 were recently shown to regulate *in vitro* condensation ^22^. Protonation of histidines with a nominative pKa of about 6.5 is regarded as a simple, dynamic, yet efficient post-translational modification that adds a positive charge upon acidification and thereby regulates ligand binding and enzymatic reactions ^36^. The pKa-values of the two histidine residues were estimated *in silico* to 6.1-6.2 in the unbound state, while the pKa of His-31 was reduced to 4.5 upon complex formation. The predicted decreased protonation likelihood of His-31 within the complex compared to the unbound state might explain the surprisingly strong binding that we observed in ITC at pH 5.6. Despite overall similar binding affinities of both ligands in ITC at pH 5.6, we observed large differences in their capacity to stabilize G3BP1-NTF2 at pH 5.1 in thermal stability measurements. G3BP1-NTF2 alone was destabilized by about 18°C relative to that at pH 7.4. This instability of NTF2 was partially compensated by ligand binding, but the stabilizing effect was increased by about 5°C for USP10 compared to Caprin-1. This effect was reduced to 2.5°C at pH 6, and abolished at neutral pH. Furthermore, our biochemical data demonstrated that G3BP1 was removed from the complex with Caprin-1 more easily at acidic pH, while the Caprin-1/FMR1 interaction was not affected (Figures 6A and 6B). In a final control experiment, exchange of Caprin-1^360-383^ by USP10^1-25^ in a chimeric Caprin-1-FGDF construct increased bound G3BP1 levels, and importantly rendered the interaction pH-independent.

Herein we have provided structural and biochemical evidence, that (i) Caprin-1^360-383^ but not USP10^1-25^ is in contact with two pH-sensitive G3BP1-NTF2 histidine residues, (ii) the resulting unique interaction interface of Caprin-1^360-383^ results in slightly lower affinity and that (iii) the stability of the G3BP1-NTF2 interaction with Caprin-1 WT but not chimeric Caprin-1-FGDF is reduced at acidic pH. Low-pH induced protonation of histidine residues may constitute a simple, evolutionarily conserved, rapid and reversible post-translational modification, regulating protein-protein interactions and thereby condensation ^36,44^. In yeast, glucose starvation induces stress and acidifies the cytoplasm, as directly shown by use of the pH-sensitive ratiometric GFP-derivative pHluorin ^51^. This acidification is characterized by the reversible transition of the fluid cytoplasm into a solid-like dormancy state ^33^. At least two proteins, the translation termination factor Sup35 and the poly-(A)-binding protein Pab1, acted as pH- and thermal stress sensors to form hydrogels at acidic pH ^11,52,53^. While acidic pH was applied *in vitro* to alter the conformational state of G3BP1, we did not find any documented evidence of acidic pH in mammalian SGs. We therefore fused the pH-sensitive GFP variant pHluorin2 to G3BP1-WT to reveal a pH-drop within SGs compared to the adjacent cytoplasm (Figure 6C and 6D) ^54^. While we are careful not to provide absolute pH-values from our measurements, we estimate that the condensates are acidified by about 0.5-1 pH-units. We speculate that the molecular origin of the observed lower pH is due to high local concentrations of RNA and/or granule-associated proteins that create this microenvironment ^8,53^. Similarly reduced pH has been observed in the nucleolus, relative to the nucleoplasm, likely due to the high concentration of nascent ribosomal RNAs therein ^55^. Our findings suggest the following regulation mechanism that supports network-theory based G3BP1-condensation models^4,20–22^: Under normal cytosolic conditions, the Caprin-1/G3BP complex stably competes with the other known NTF2-hub binding proteins such as USP10 and Ubap2/Ubap2L ^20–22^. Upon stress, Caprin-1/G3BP1 coalesce into SGs mediated by multivalent higher-order RNA-interactions, resulting in reduced pH. In the resulting acidified microenvironment, the Caprin-1/G3BP1-NTF2 interaction is destabilized such that USP10 gains a competitive advantage for NTF2-binding, thus allowing for rapid recycling of G3BP1 and ultimately for condensate disassembly upon stress release.

In this manner, reduced pH in SGs could regulate and fine-tune the assembly and disassembly of biomolecular condensates, not only by conformational changes within full-length G3BP1 (and likely other interactions), but also via a NTF2-mediated pH-sensitive SLiM-selection mechanism. The various roles of Caprin-1 in cancer as a cell cycle regulator and in neurodegenerative disease as potential contributor to altered RNA metabolism and also the roles of G3BP in viral infections, cancer and neurodegeneration disease highlight the importance of this pH-regulated interaction. This work thus uncovers new targets for the potential treatment of diseases characterized by dysregulation of SGs ^7,56–58^.

## Materials and methods

### Heterologous protein production

G3BP1-NTF2 (residues 1-139) was bacterially expressed and purified as previously described ^31^. Briefly, the G3BP1-NTF2 dimer was affinity-purified (Immobilized Metal Affinity Chromatography (IMAC), HisTrap FF, GE Healthcare) and isolated from a Superdex 75 size exclusion chromatography (S75, SEC, GE Healthcare). After TEV cleavage (PSF, KI, Stockholm), G3BP1-NTF2 was collected as IMAC flow-through and re-applied on the same S75 column. For crystallization, the Caprin-1-derived peptide comprising residues 356-386 (RQRVQDLMAQMQGPYNFIQDSMLDFENQTLD) was dissolved in dH2O and added to G3BP1-NTF2 in 5 times molar excess. The G3BP1-NTF2:Caprin-1^356-386^ mix was dialyzed against 20 mM HEPES, 300 mM NaCl, 10% glycerol, 1 mM TCEP, pH 7.5 (HEPES-buffer TCEP) and concentrated to a total absorbance of 8.5 using ultrafiltration with a MW cutoff of 3 kDa. DNA fragments encoding Caprin-1^356-386^ and USP10^1-28^ were ligated to the 3’ end of fragment encoding GFP containing N-terminal twin-Strep-tag-II and TEV-cleavage sites on pET21d-expression vectors (in-house, GFP-USP10^1-28^ and GFP-Caprin-1^356-386^). These modified vectors and expression constructs were obtained by sequence and ligation independent cloning (SLIC) and validated by DNA sequencing (Eurofins Genomics) ^59,60^. GFP-USP10^1-28^ and GFP-Caprin-1^356-386^ were expressed in *E. coli* BL21 T7 Express. Bacterial pellets were lysed in HEPES-buffer comprising 0.4 g/L lysozyme, 0.05 g/L DNase, 1 mM PMSF and 1 tablet/50 mL Roche EDTA-free protease inhibitor cocktail by a mechanical pressure cell (Homogenising Systems Ltd). STII-GFP-USP10^1-28^ and STII-GFP-Caprin-1^356-386^ were affinity-purified using Strep-Tactin columns (IBA). Tags were cleaved by incubation with TEV-protease in HEPES buffer containing 1 mM EDTA and 1 mM DTT. Cleaved proteins were isolated from a Superdex 75 column (GE Healthcare) equilibrated in ITC-buffer (25 mM HEPES, 150 mM NaCl, 10 mM MgCl_2_, 10% glycerol, pH 7.5).

### Crystal structure determination

Crystals were refined around the initial hit condition comprising 0.2 M NaCl, 2M sodium chloride, 0.1 M Tris pH 8.0, 20% (w/V) PEG 4000 from the Proplex crystallization screen (Molecular Dimensions), and setup in 96-well sitting drop iQ plates using the Mosquito LCP robot (TTP Labtech). Crystals were cryo-protected by soaking in mother liquor supplemented with 30% (w/V) glucose, and flash-frozen in liquid nitrogen. X-ray data were collected at beamline BL14-1 at the BESSY synchrotron radiation facility (Berlin, Germany) ^61^. Diffraction data were collected to a resolution of about 1.9 Å and processed using the XDS/XDSAPP program package ^62,63^ (Table S1). The molecular replacement software Phaser placed four NTF2 monomers into the asymmetric unit (asu) that were manually extended to comprise six NTF2 molecules as evident from the electron density in Coot ^64,65^. The R and Rfree values of the refined manually extended MR solution were 28.6 and 34%, respectively. The electron density clearly revealed additional density (discovery map) to build 20 residues of the co-crystallized 31-AA long Caprin-1^356-386^ peptide bound to chain D, and the final model was refined further to R and R_free_ values of 20.8 and 25.2%, respectively. Coot and Phenix were used to manually re-build and automatically refine the model, comprising non-crystallographic restraints, individual isotropic B-factors and automatically determined TLS groups ^64,66^. Residue-level map-to-model correlation plots were obtained using Phenix and R tidyverse ^66,67^. Structural analysis was performed using PDBePISA, PyMol, Coot, PIC, and ROSIE web servers ^45,64,68–73^. Residue-level interaction summary plots and associated PyMol scripts were generated using R and the tidyverse package to analyze interaction tables obtained from Molprobity ^67,74–76^.

### Biophysical measurements

#### Isothermal titration calorimetry (ITC)

Buffers of GFP-USP10^1-28^ and GFP-Caprin-1^356-386^ as well as G3BP1-NTF2 were exchanged to ITC-buffer (25 mM HEPES, 150 mM NaCl, 10 mM MgCl_2_, 10% glycerol, pH 7.5) by S75-SEC. ITC measurements were performed using an ITC200 calorimeter (GE Healthcare). The other buffers used to collect biophysical ITC or nanoDSF data at different pH-values were 25 mM MES pH 6.1 or pH 5.6 as well as 25 mM Na-acetate pH 5.1, 150 mM NaCl, 10 mM MgCl_2_, 10% glycerol (V/V). The cell temperature was set to 25°C and the syringe stirring speed to 750-1000 rpm. Before each experiment, G3BP1-NTF2 and GFP-USP10^1-28^ or GFP-Caprin-1^356-386^ were loaded into cell and syringe at concentrations of 10-25 μM and 150-500 μM, respectively. Data and binding parameters were preanalyzed using the MicroCal PeakITC software (Malvern). Final analysis and global fits were performed using NITPIC and Sedphat, respectively ^77,78^.

#### Prometheus nanoDSF

Prometheus nanoDSF measurements were performed in duplicate by loading NTF2 samples at absorbance values of 0.2 into high-sensitivity capillaries following the manufacturer’s protocols (Nanotemper). Peptide ligands were dissolved in DMSO and added to NTF2 to yield final DMSO concentrations of 4% (V/V). The DMSO concentration was kept constant at 4% during the initial titration experiments. Buffers were identical to the ITC measurements. All data were combined, tidied and visualized applying ggplot2 and the tidyverse packages in Rstudio ^67,75,76^.

#### Bio-Layer Interferometry (BLI)

All BLI measurements were performed on an Octet RED instrument (ForteBio). GFP-USP10^1-28^ and GFP-Caprin-1^356-386^ proteins were immobilized covalently on the surface of Amine Reactive Second-Generation (AR2G) Biosensors following the manufacturer’s protocols (ForteBio). Surfaces were regenerated applying cycles comprising three 5 s-dips in 10 mM HCl and HEPES buffer after and before each titration experiment. For the titration experiments, four of the eight sensors in each sensor set were used as reference and treated only with sulfo-NHS/EDC and ethanolamine, but not loaded with GFP-USP10^1-28^ and GFP-Caprin-1^356-386^. The stability of the sensor surfaces was monitored by including 500 nM NTF2_2_-samples in each experiment. In a second set of comparative experiments, GFP-USP10^1-28^ and GFP-Caprin-1^356-386^ wild-type and mutant proteins were loaded on two and one sensor surfaces, respectively. One of the remaining sensors was used as blank, and the other as additional GFP-USP10^1-28^ wild-type surface. In this second set of experiments, NTF2_2_ was applied at the same concentration for all sensors and measured in parallel. Data were collected in 25/25 mM HEPES/MOPS pH 7.4, 1% glycerol (V/V), 0.05% Tween-20 (V/V), 150 mM NaCl. Raw data were pre-processed by subtracting the reference from the sample surfaces and applying a Savitzky-Golay filter as implemented in the Octet RED analysis software.

### Cell lines and mutant generation

#### Applying CRISPR-Cas9 to obtain ΔCaprin-1 cell line

U2OS-WT cells were plated and transfected with the pCas9-Guide (Origene GE100002) constructs using Lipofectamine 2000 overnight, allowed to recover for ≥2 days, and thereafter were reseeded and immuno-stained for Caprin-1 and G3BP1. Cultures with less than 5% KO cells were first “pool cloned” to enrich for KOs by plating at 5–10 cells per well in 24 well plates, allowing the cells to grow to >50% confluency before reseeding on coverslips in a 24-well plate for screening. When the cells on coverslips reached 80% confluency, cells were fixed and stained for Caprin-1 and G3BP1. Samples showing desired KO >5% were subcloned by limiting dilution. To identify Cas9-induced mutations in the Caprin-1 coding sequence, genomic DNA was isolated using Trizol reagent (Thermo Fisher). Amplification of the genomic DNA was performed using a primer set spanning Exon 2 of genomic Caprin-1. Primer 1, 5’-GCCCCGTCCGTCTCCTG-3’ (forward) and primer 2, 5’-AGGAGGGGACCTAGTAACGCTC-3’ (reverse). Genomic DNA PCR was performed with DreamTaq PCR Master Mix (Thermo Fisher). DNA was initially denatured at 95°C for 1 min, followed by denaturation at 95°C for 30 s, annealing at 60°C for 30 s, and extension at 72°C for 1 min for 30 cycles. Final extension was done at 72°C for 5 min. PCR products were gel purified and directly cloned into pCR.2.1 vector (Thermo Fisher). Individual clones were propagated, plasmids isolated and sequenced.

#### Other cell lines and transient transfections

U2OS-WT, ΔΔG3BP1/2 U2OS ^2^, ΔCaprin-1 U2OS cells were maintained at 5.0 % CO_2_ in DMEM containing, 10% fetal bovine serum, 100 U/mL penicillin and 100 μg/mL streptomycin. Stable ΔΔG3BP1/2 U2OS-derived cell lines constitutively expressing fluorescently tagged proteins (G3BP1-WT and mutants) were made as described in detail elsewhere ^79^ by transfecting pEGFP-C1-G3BP1 and mutants into ΔΔG3BP1/2 U2OS cells, selecting with 100 μg/ml G418 (Gibco) for 7-10 days, and screened using fluorescence microscopy and Western blotting. For transient transfections cells were grown to 70-80% confluency and transfected using Lipofectamine 2000 (Invitrogen) following the manufacturer’s instructions and processed after 24 h.

### Molecular cloning

#### Creation of EGFP-G3BP1 and AcGFP-Caprin-1 mutants

The 5’ phosphorylated primers (Table S3) were mixed with 20 ng of pEGFP-C1-G3BP1-WT or pAcGFP-C1-Caprin-1-WT according to protocol with Phusion DNA polymerase (Thermo Fisher) at a final volume of 20 μL. The mixture was denatured at 98°C for 45 s, followed by 28 cycles of the following: 98°C for 15 s, 61-66°C (depending on primer set) for 15 s, 72°C for 4 min, with a final extension step of 72°C for 12 min. The PCR mixture was incubated with one-unit DpnI (NEB) for 1h at 37°C in a final volume of 40 uL to remove *E. coli* derived template DNA, followed by heat inactivation at 80°C for 20 min. 50 ng of the PCR product was ligated with T4 DNA ligase (NEB) in a final volume of 10 μL overnight at 4°C. 5 μL of the ligation mix was used for chemical transformation into high-efficiency 10 beta *E. coli* (NEB).

#### Creation of pAC-GFP-Caprin-1^Δ360-383^-USP10^1-25^ (GFP-Caprin-1-FGDF chimera)

This plasmid was created using NEBuilder HiFi DNA assembly protocol (NEB). In short, backbone pAc-GFP-Caprin-1^Δ360-383^ was created via Phusion PCR protocol (see above) with primers (Table S3) containing a 20 nt overhang to the USP10^1-25^ insert. This USP10^1-25^ insert was also amplified with primers (Table S3) containing a 20 nt overhang to the pAc-GFP-Caprin-1^Δ360-383^ backbone via a Phusion PCR protocol (see above) based on a pEGFP-C1-USP10^1-40^-wt template ^28^. Both PCR products were mixed according to the manufacturer instruction and 10 beta *E. coli* (NEB) were chemically transformed. Plasmids were isolated and sequenced.

### Immunoprecipitation and immunoblotting

#### Standard procedure

60 mm dishes of 70-80%-confluent of ΔCaprin-1 U2OS cells were transiently transfected for 24 h, washed with PBS, and scrape-harvested at 4°C into EE lysis buffer (50 mM HEPES, 150 mM NaCl, 2.5 mM EGTA, 0.5% NP40, 10% glycerol, 5 mM EDTA, 1 mM DTT, HALT protease inhibitors (Thermo Fisher). ΔΔG3BP1/2 U2OS stably expressing GFP-G3BP1-wt and mutated versions (G418 selected) were grown in 6 well plates until 70-80% confluency washed with PBS and scrape harvested at 4°C into EE lysis buffer. Lysates were rotated for 10 min at 4°C, sonicated for 8 min on ice, cleared by centrifugation (10.000 g, 10 min, 4°C), and incubated with anti-GFP beads for 1 h with continuous rotation at 4°C. Anti-GFP beads were produced by expressing the GFP nanobody “Enhancer” ^80^ in *E.coli,* purified by size-exclusion, and coupled to cyanogen bromide-activated sepharose (Sigma-Aldrich) or commercial GFP-Trap agarose (Chromtek). Beads were washed in EE lysis buffer and eluted directly into 2x NuPAGE LDS sample buffer with 100 mM DTT and denatured for 5 min at 95°C. Proteins were resolved in 4-12% Bis-Tris NuPAGE gels (Thermo Fisher Scientific) and transferred to nitrocellulose membranes using the Trans-Blot Turbo Transfer system (Bio-Rad) and blotted using standard procedures. Chemiluminescence was detected using SuperSignal West Pico substrate (Thermo Fisher).

#### pH dependent immunoprecipitation

20×10^6^ ΔCaprin-1 U2OS cells stably expressing GFP-Caprin-1-WT or GFP-Caprin-1-FGDF were lysed in 1.5 mL (50 mM HEPES, 150 mM NaCl, 2.5 mM EGTA, 0.5% NP40, 10% glycerol, 1 mM DTT, HALT protease inhibitors (Thermo Fisher), 50 μg/mL RNase A (NEB). Lysates were rotated for 10 min at 4°C, sonicated for 8 min on ice, cleared by centrifugation (10000 g, 10 min, 4°C), and incubated with GFP-Trap agarose beads (ChromoTek) for 1-2 h with continuous rotation at 4°C. Then beads were pelleted at 2000 g for 1 min, resuspended and equally divided into four fresh 1.5 mL tubes. Aliquoted beads were pelleted and drained from supernatants. Drained beads were suspended and rotated in 1 mL of pH specific buffers for 10 min (150 mM NaCl, 10 mM MgCl2, 10% glycerol, 0.5% NP40 in 25 mM MES pH 5.6 / pH 6.1 / 50 mM HEPES pH 7 / 25 mM HEPES, 25 mM MOPS pH 7.4). Beads were pelleted and pH specific buffers were removed, subsequently washed twice with the same pH specific buffer, and twice with pH 7.0 buffer. The last buffer was completely removed from the beads, and complexes were eluted with 2x NuPAGE LDS sample buffer with 100 mM DTT, boiled for 5 min at 95°C. Samples were analyzed by western blotting as above.

Western blots were quantified (BioRad ImageLab 6.0.1) and analyzed in GraphPad Prism. Data shown are mean ± SEM and are analyzed using the unpaired t test. ns; non-significant, *, P < 0.05; **, P < 0.01.

### Immunofluorescence analysis

#### Standard procedure

Cells were fixed and processed for fluorescence microscopy as described previously ^81^. Briefly, cells were grown on glass coverslips, stressed as indicated, fixed using 3.7 % formaldehyde (Sigma) in PBS for 10 min, followed by 5 min post-fixation/permeabilization in ice-cold methanol. Cells were blocked for 1 h in 5% horse serum/PBS, and primary and secondary incubations performed in blocking buffer for 1 h. All secondary antibodies were raised in donkey against either mouse, rabbit or goat and tagged with either Alexa Fluor 488, 568 or 647 (Thermo Fisher). Following washes with PBS, cells were mounted in polyvinyl mounting media and viewed using a Zeiss Axiovert 200M with a 63X Plan Apochromat objective lens (NA 1.4) and illuminated with a mercury lamp and standard filters for DAPI (Filter Set 49: 335-383 / 395 / 420-470), Alexa Fluor 488 (Filter Set 10: 450-490 / 510 / 515-565), Alexa Fluor 568 (Filter Set 20: 540-552 / 560 / 575-640), Alexa Fluor 647 (Filter Set 50: 640/30 / 660 / 690/50). Images were captured using an AxioCam ICc5 digital camera (Zeiss) with the manufacturer’s software, and raw TIF files were imported into Adobe Photoshop. Identical adjustments in brightness and contrast were applied to all images in each experiment.

#### High content microscopy

For high content microscopy, ΔΔG3BP1/2 U2OS cells stably expressing GFP-G3BP1-WT and mutants were plated on a μ-Plate 24 Well Black plates ibiTreat (IBIDI) and grown for additional 24 h. Cells were left unstressed or stressed with 200 μM sodium arsenite for 1 h, then fixed and stained as described above. Images were recorded with a Molecular Devices ImageXpress Micro microscope, equipped with a 20x objective, and illuminated with a mercury lamp and standard filters for DAPI, FITC, Cy3 and Cy5. Images were captured using a four-megapixel sCMOS digital camera with the manufacturer’s software MetaXpress, and raw TIF files were analyzed with CellProfiler ImageJ and Rstudio. False positives were mainly observed for mock treated cells due to oversaturated areas that led to artificially created SGs around the periplasmic region. We estimate the false positive rate to be lower for stressed cells since the parameters of the algorithm were tuned to detect “speckle” signals in the cytoplasm above background as SGs.

#### Ratiometric pHluorin2-G3BP1-wt measurement

ΔΔG3BP1/2 U2OS cells were transfected with pHluorin2-G3BP1-wt for 48 h using Lipofectamine 2000, then selected with 100 μg/mL G418 for 7-10 days. Selected cells were plated on μ-Plate 24 Well Black plates ibiTreat (IBIDI) and grown for additional 24 h. Cells were then left untreated or treated with 200 μM sodium arsenite diluted in complete media or 20 μM clotrimazole diluted in serum-free media for 30 min then viewed with a Supercontinuum Confocal Leica TCS SP5 X, equipped with a pulsed white light laser, 405 nm violet diode laser, a Leica HCX PL Apo 63x/1.40 oil objective and a heated chamber set on 37°C. Live cell images were recorded sequentially with the following settings to avoid saturated signals. Pinhole was set to 2 airy. Firstly: Excitation at 485 nm with 25% intensity of the white laser light (WLL), PMT1 (photomultiplier tube) detector was set to Gain 900 V and Offset −1.0 V, emission window was set from 505-515 nm. Secondly: Excitation at 405 nm of the violet diode laser with 25% intensity, PMT1 detector was set to Gain 900V and Offset - 1.0 V, emission window was set from 505-515 nm. Quantification was performed on 4 images with 3-4 cells and multiple SGs/cell, which totaled around 40 or more SGs/per experiment. The area of the region of interest was set equal in SGs and in adjacent cytoplasm and analyzed. Then the mean pixel values of channel 485 nm and 405 nm were measured and 405/485 nm ratio was calculated and plotted ^48^.

## Contributions, Acknowledgements, Funding and data availability

### Authors contribution

MDP conceived the study. MDP and TS designed all experiments and analyzed the data. AA and GMM were responsible for funding acquisition and oversaw the overall project as PIs. TS and MDP wrote the initial draft that was revised together with AA and GMM. All authors contributed to the interpretation of the results and the final version. / by methods: X-ray crystallography: TS; biophysics: TS with support from JSF, AO, JN, KAG and PÅN. Cell biology, biochemistry and immunofluorescence: MDP, LW, TJCT with support from LH, AMM.

## Acknowledgements

Part of this work was facilitated by the Protein Science Facility at Karolinska Institutet/SciLifeLab (http://ki.se/psf), and we thank Martin Moche and Tomas Nyman for excellent technical support for X-ray crystallography and ITC. Crystallographic computations and data handling were enabled through the Presto installation provided by the Swedish National Infrastructure for Computing (SNIC) partially funded by the Swedish Research Council through grant agreement no. SNIC 2020/5-368. We acknowledge excellent technical assistance and instrument access to surface plasmon resonance, BLI and nanoDSF through Aman Mebrahtu (KTH Stockholm) and Yasmin Andersson (SciLifeLab Stockholm). We thank HZB for the allocation of synchrotron radiation beamtime. This project has received funding from the European Union’s Horizon 2020 research and innovation programme under grant agreement No 730872. We also thank Tatyana Sandalova and Francesca Pennacchietti (SciLifeLab Stockholm) for critically reviewing the manuscript.

## Data availability

Raw data and analysis scripts and will be made publicly available upon peer-reviewed publication (via DataDryad/Github). The crystal structure was deposited with PDB-ID 6TA7.

## Abbreviations used in this study

BLI: Bio-Layer Interferometry
BMCs: Bio-molecular condensates
Caprin-1: Cell cycle-associated protein 1
co-IP: Co-immunoprecipitation
ECDF: Empirical cumulative distribution function
FMR1/FMRP: Fragile X mental retardation protein
G3BP1: Ras GTPase-activating protein-binding protein 1
IP: Immunoprecipitation
ITC: Isothermal titration calorimetry
LCR /IDR: Low complexity or intrinsically disordered regions
nanoDSF: Nano differential scanning fluorimetry
NTF2: Nuclear transport factor 2 like domain
RGG: Arginine-Glycine-Glycine motif
SA: Sodium arsenite
SEC: Size exclusion chromatography
SFV: Semliki Forest virus
SG: Stress granule
SLiM: short linear (peptide) motif
USP10: Ubiquitin carboxyl-terminal hydrolase 10
WT: Wild type

## Supplemental material

### Supplement Figures

**Figure S1.**
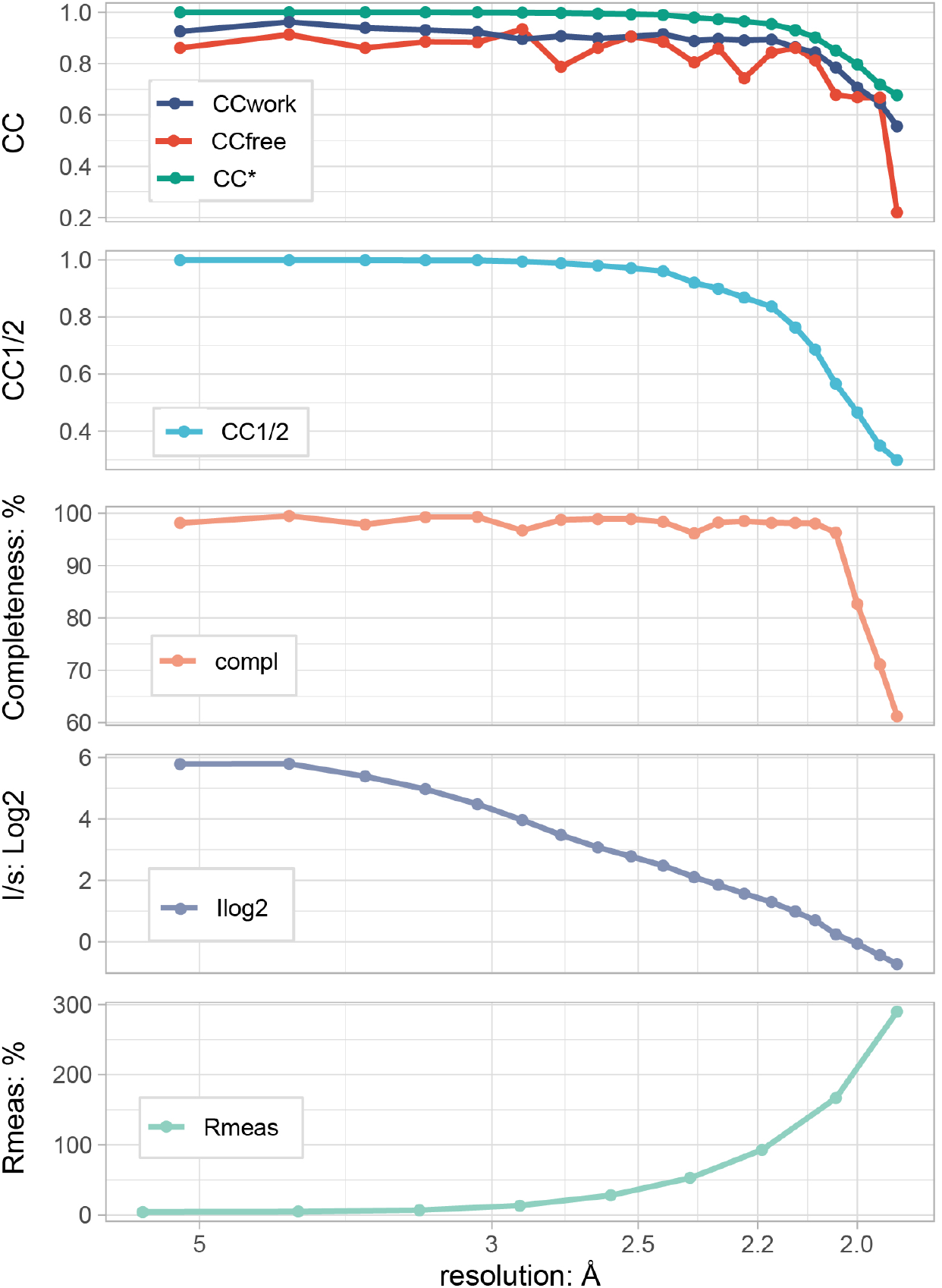
Crystallographic quality indicators for PDB structure 6TA7. Cross correlation (CC) and other quality indicators^1^: *CC1/2:* CC between intensity estimates from half data sets; *CCstar:* Calculated from CC1/2. Indicator useful for comparing data and model quality. *CCwork and CCfree:* the standard and cross validated correlations of the experimental intensities, with the intensities calculated from the refined molecular model. *I/σ:* Average signal-to-noise ratio of individual observations, given on log2-scale. Data fall below I/ *σ* < 2 at a resolution of about 2.1 Å. *Rmeas:* Multiplicity independent replacement of Rmerge and Rsym; Useful for assessing space group symmetry and isomorphism of multiple data sets; Should play no role in determining resolution cutoff. *Completeness:* Percentage of measured unique reflections out of the expected ones.

**Figure S2.**
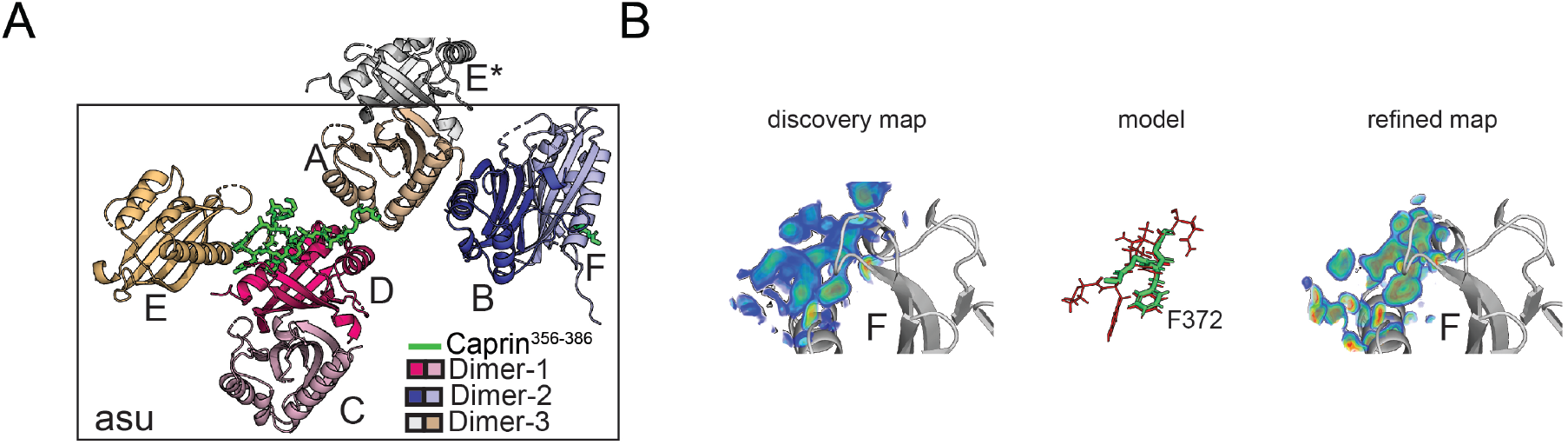
The asymmetric unit contains three NTF2 dimers and two 20AA-long and 4AA-short fragments of Caprin-1^356-386^ bound to chains D and F. (A) The asymmetric unit of the G3BP1-NTF2/ Caprin-1^356-386^ crystal structure contains three nonidentical NTF2-dimers created by chains C/D, A/E and B/F. The 20AA-long and 4AA-short Caprin-1 fragments bound to NTF2 chains D and F, respectively. (B) The discovery and refined maps as well as the built model of Phe-372 and few additional backbone atoms of Caprin-1^356-386^ bound to chain F are visualized similarly to the long Caprin-1^356-386^ fragment in Figure 1B.

**Figure S3.**
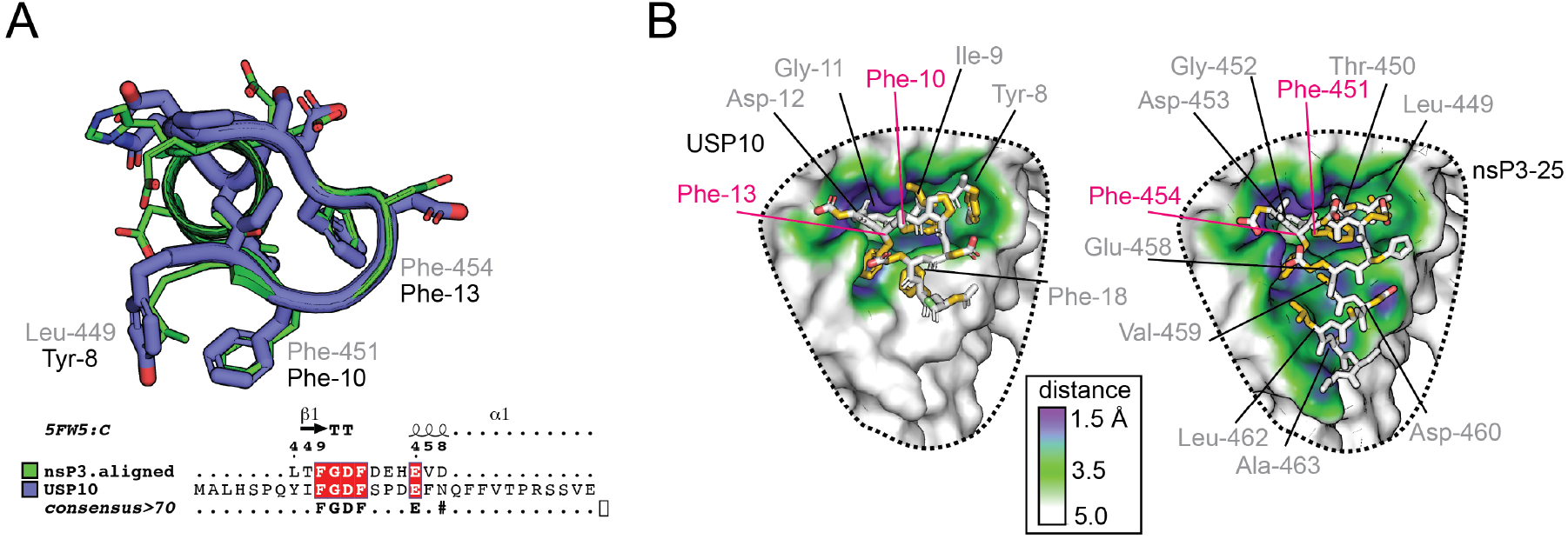
Molecular model for USP10 binding to NTF2. (A) The structure of USP10 was predicted based on nsP3^449–473^ as template using Phyre, and manually adapted in Coot to include Tyr-8 that was not modeled and then refined using Rosetta FlexPepDock. The aligned sequence was visualized using ESPript. ^2^ (B) The putative binding interface of the docked USP10 was visualized similarly to Figure 3.

**Figure S4:**
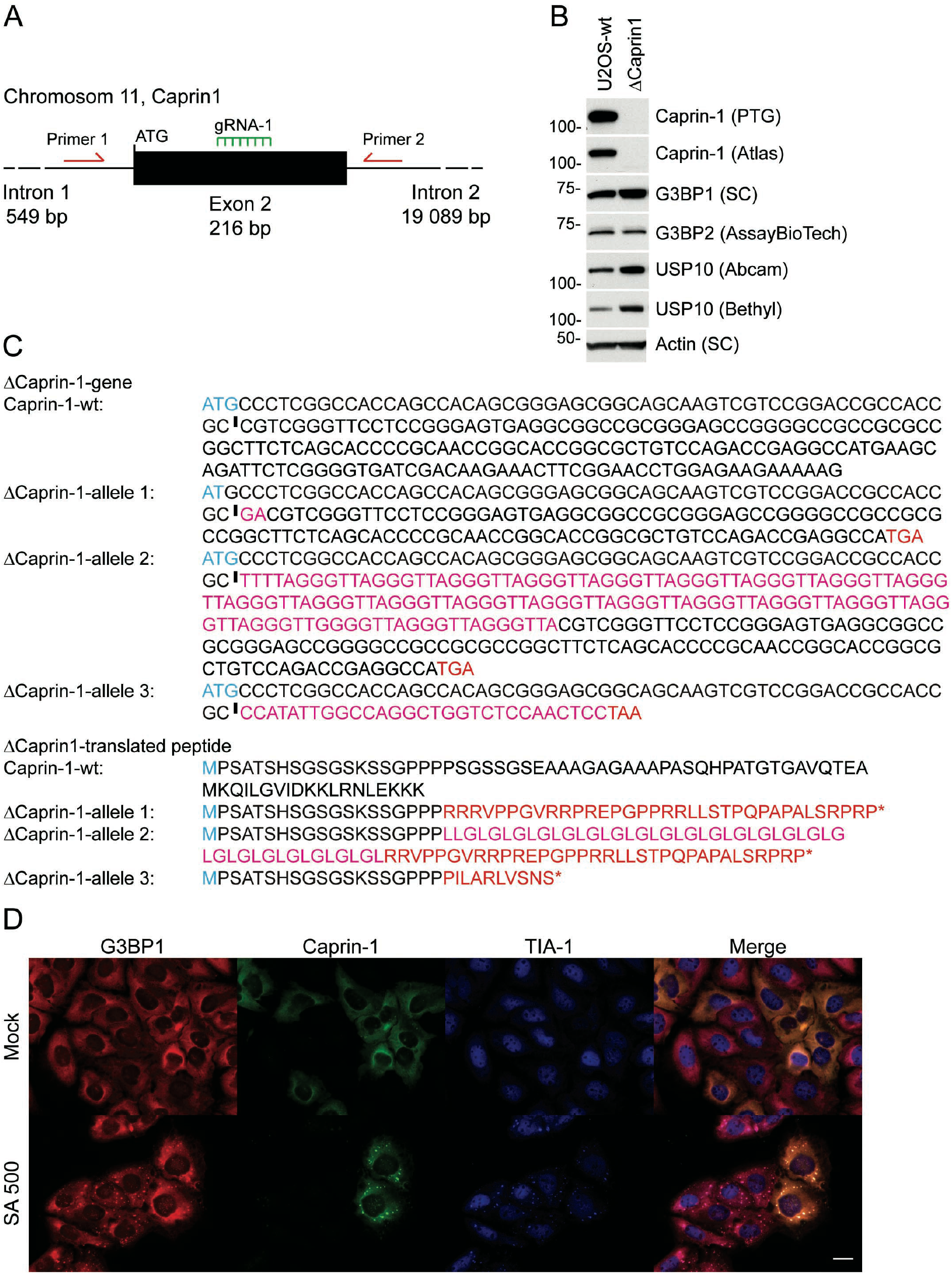
Evaluation of Caprin-1 KO cells and Creation of U2OS Caprin-1 knockout cells using CRISPR/Cas9 technology. (A) CRISPR/Cas9 mediated deletion of Caprin-1 utilizing guide RNA located within the second exon of Caprin-1. Green line indicates location of gRNA and red arrows denote primers used for genotyping and sequencing. (B) Western blotting confirming loss of Caprin-1 expression with two different commercial antibodies, while direct interaction partners G3BP1 and G3BP2 are unaffected, interaction competitor USP10 is affected. Actin serves as loading control. (C) Genotype of U2OS ΔCaprin-1 cells. Gene sequences showing Cas9-induced insertions (magenta); initiator ATG appears blue, black vertical line indicates Cas9 double-strand break, inframe premature stop appears in red. Predicted protein products of native gene are aligned above the three mutant alleles, with predicted frameshifted aa and premature stop codons (red asterisks). (D) U2OS-WT and ΔCaprin-1 cells, co-cultured, treated as indicated and stained for G3BP1 (red), Caprin-1 (green), and TIA-1 (blue). Bar: 20 μm

**Figure S5.**
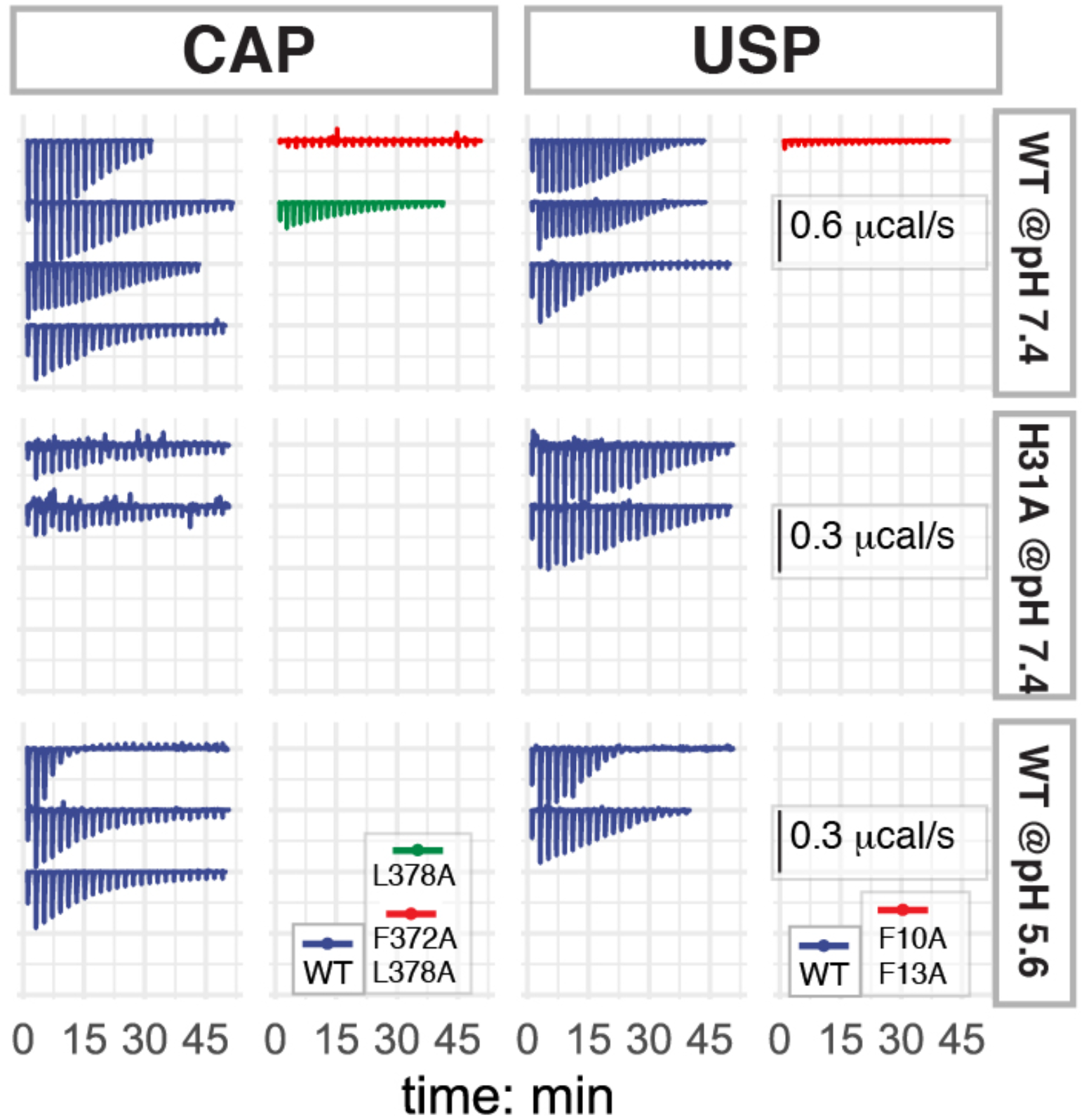
ITC thermograms. Baseline-corrected thermograms from which the integrated binding heats were derived. Thermograms are colored as in Figure 2. The measurement group comprising NTF2-wildtype at pH 7.4 was performed at approximately double concentrations compared to the other two measurement groups, as apparent from the double-scaled raw data.

**Figure S6.**
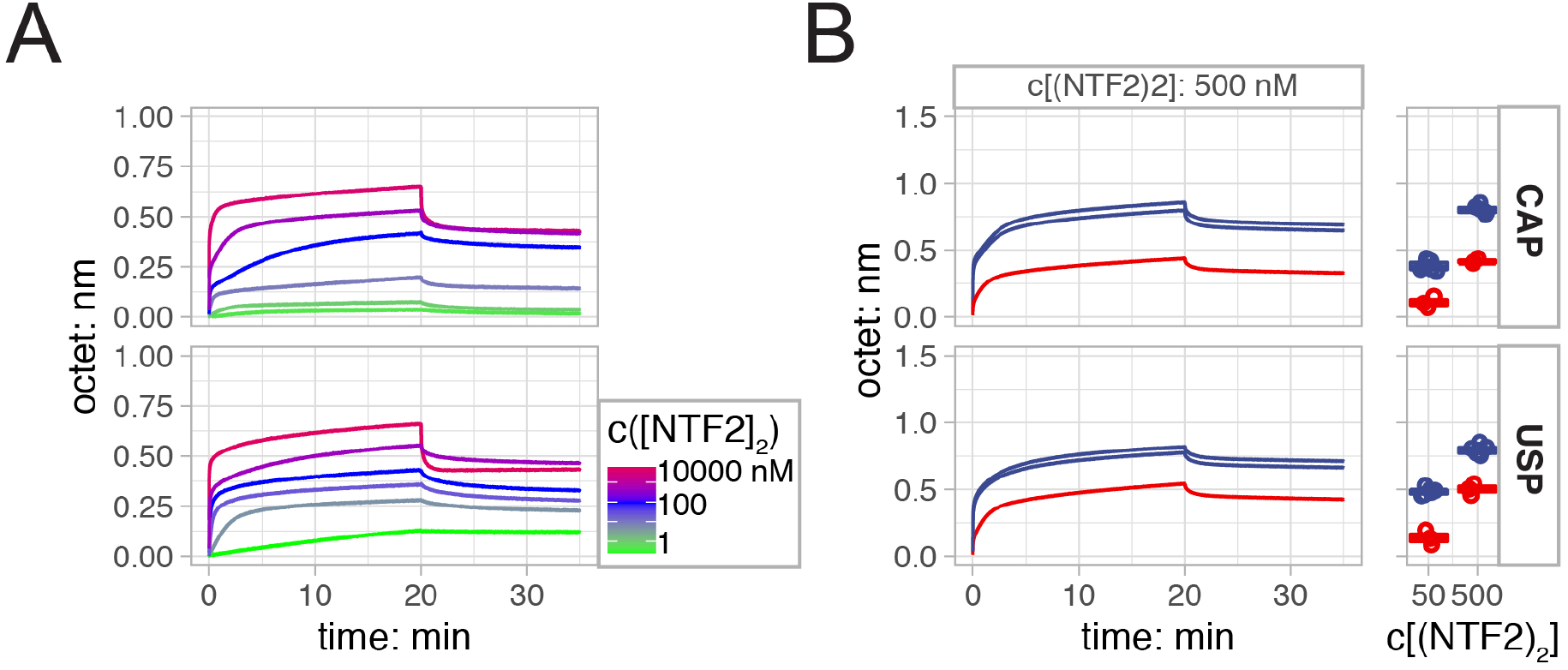
NTF2 bound with higher apparent affinity to surface-immobilized GFP-USP10 compared to GFP-Caprin-1. (A) In BLI binding of the NTF2 dimer (NTF2_2_) to surface-immobilized GFP-USP10^1-28^ and GFP-Caprin^356-386^ yielded very fast association and very slow dissociation phases for both interactions, most likely due to electrostatic attraction of NTF2 to the sensor surface and the bivalent nature of the analyte. Pseudo-equilibrium binding levels were derived from the final timepoints of the association phases of each titration. Representative curves from in total 32 and 40 curves for CAP and USP, respectively. (B) *left panel:* Kinetic binding curves measured at NTF2-concentration of 500 nM. *right panel:* Boxplots of binding response levels at NTF2-concentrations of 50 and 500 nM from for three measurement repeats of each set. Binding levels of 500 nM NTF2 to the double mutants reached approximately the binding levels of wild-type NTF2 at 50 nM, indicating that binding of NTF2 to the mutants was estimated to be about 10 times weaker.

**Figure S7.**
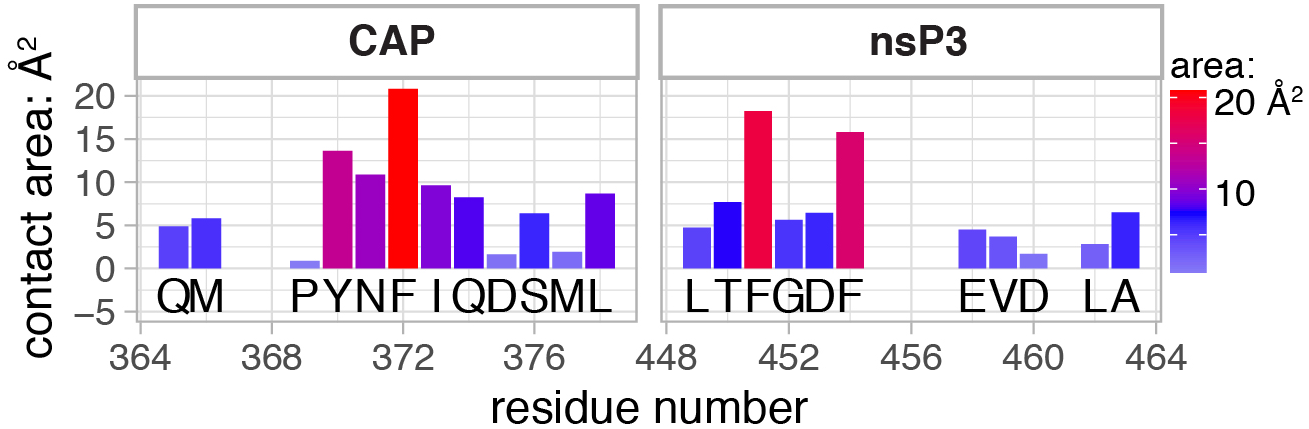
Aromatic anchor residues highlighted by interface areas. Residue-level contact interface areas integrated from the Molprobity output tables highlighted key aromatic anchors of Caprin-1 and nsP3, as well as contact hot spots.

**Figure S8.**
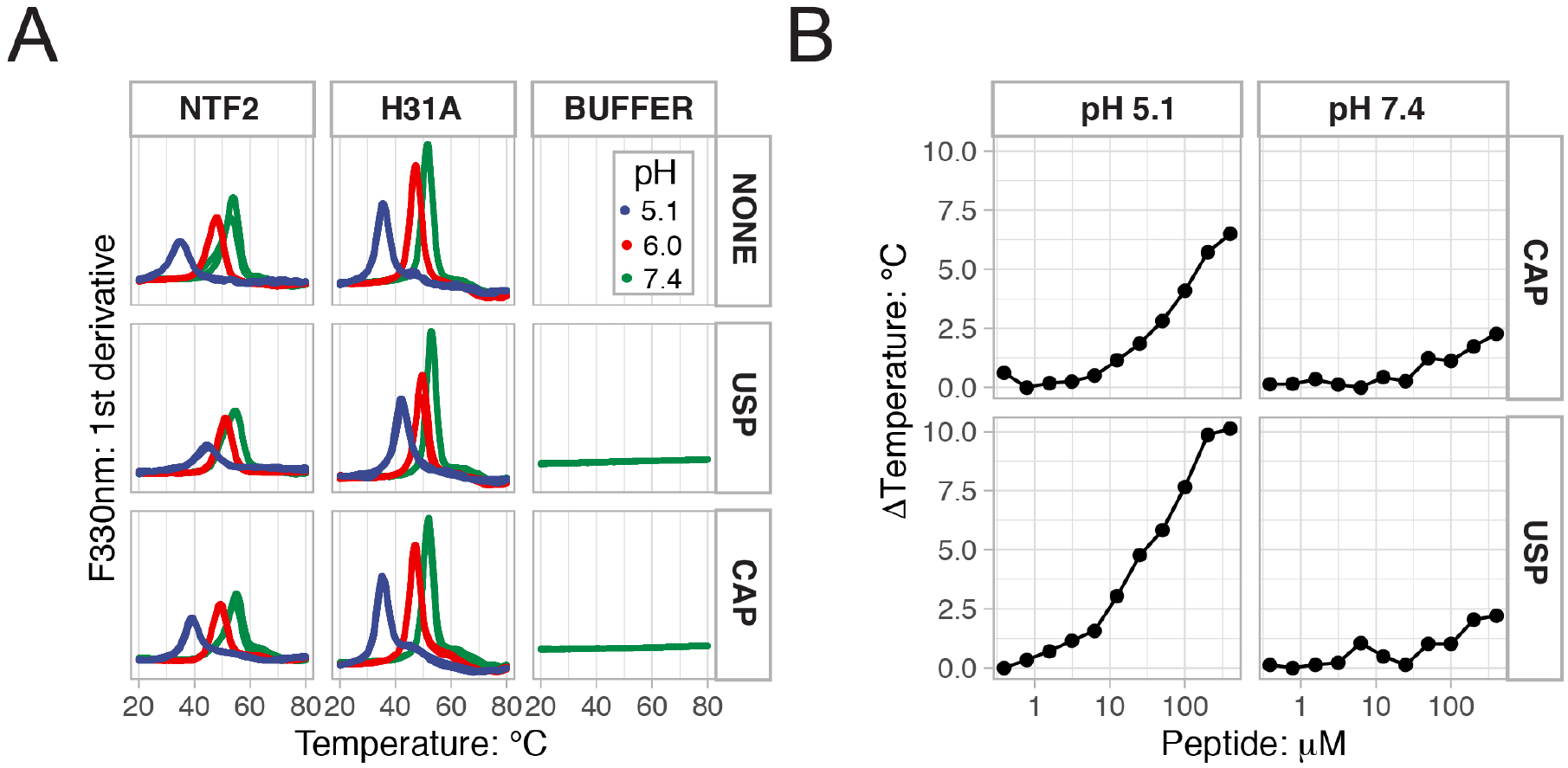
nanoDSF demonstrated decreased NTF2 stability at acidic pH, that is compensated by USP and CAP ligand binding. In thermal unfolding experiments using the Prometheus nanoDSF system, protein samples are heated at a constant rate and thermal melting is monitored as a change in fluorescence of intrinsic aromatic residues at high resolution, as well as changes in light scattering due to aggregate formation (https://nanotempertech.com/). (A) Thermal unfolding of G3BP1-wildtype (top panel) displayed midpoint transition temperatures (Tm, first derivative peak maximum) of 53, 48 and 35°C when measured at buffer pH-values of 7.4, 6.0 and 5.1, respectively. Similar values were obtained for the H31A mutant. Incubation of G3BP1 with about 10 times molar excess of peptide ligands shifted the peaks to higher temperatures. The peptide alone did not display any midpoint transition temperature. (B) Initial peptide titrations demonstrated concentration-dependent stabilization of NTF2. The stabilization assays summarized in Figure 4D were performed at NTF2 and peptide concentrations of 20 and 200 μM, respectively.

**Figure S9.**
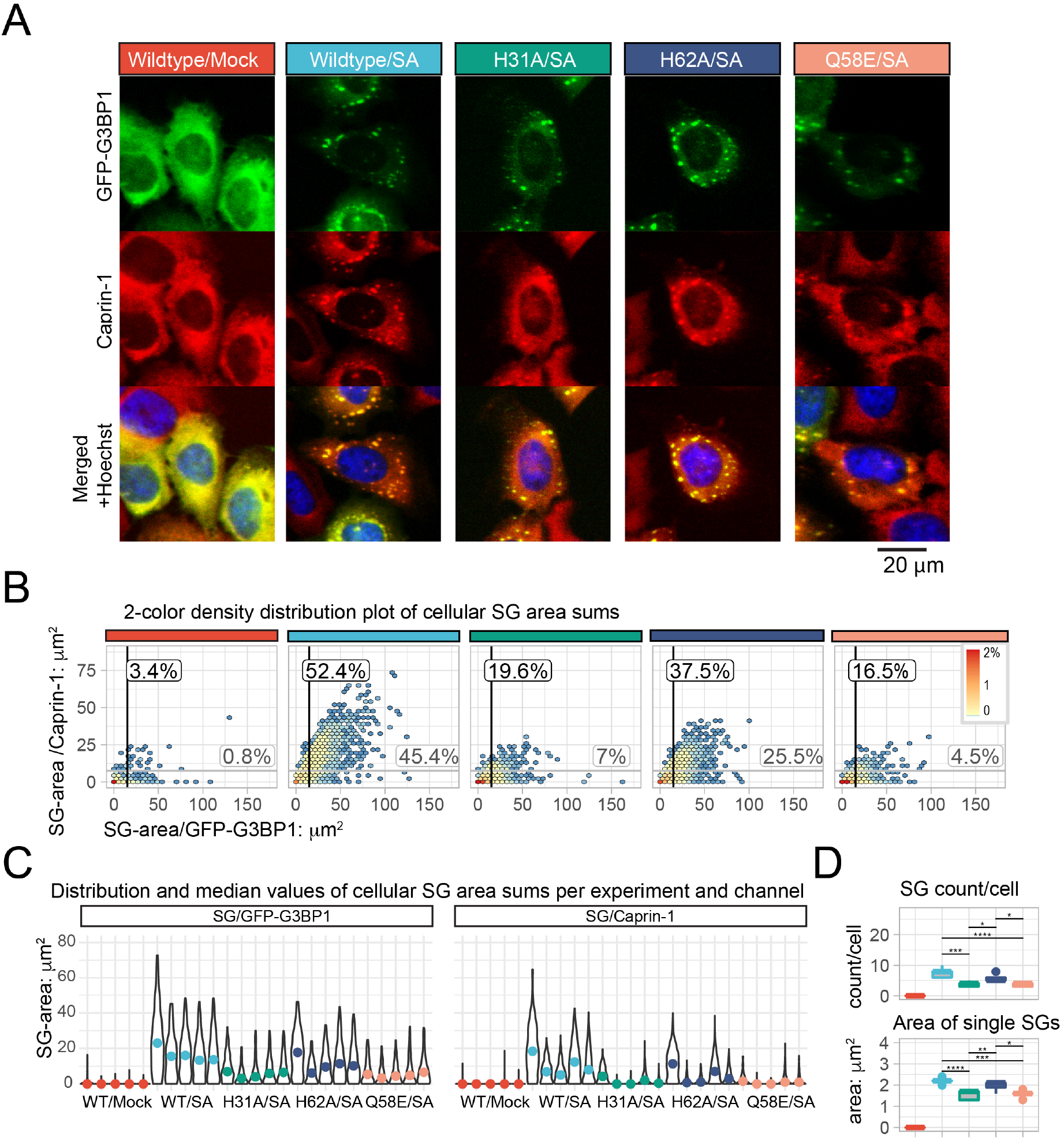
Representative high-content microscopy images and additional data statistics. (A) Representative high-content microscopy images of the green (GFP-G3BP1), red (endogenous Caprin-1) and merged (plus blue Hoechst staining of nuclei) channels. (B) The SG area sums per cell were binned and the counts visualized as cell densities on the twodimensional channel grid. SG areas are given in μm^2^. Cells with increased GFP-G3BP1 and Caprin-1 SG area sums are shifted to the right and upwards within the two-dimensional grid. Data were combined from all five experiments and gates were set at 15 and 7.5 μm^2^ in the green and red channels, respectively, yielding percentages of 52 and 42% for G3BP1-WT cells, compared to 38 and 26%, 20 and 7% as well as 17 and 5% for G3BP1-H62A, G3BP1-H31A as well as G3BP1-Q58E, respectively. (C) The distribution of cellular SG-area sums is visualized as violin plots for each cell population and experiment, and separated into two panels for the GFP-G3BP1 and Caprin-1 channels. Distributions are scaled such that the maximum count of each population has the largest width, i.e. 85% of mock-treated cells have SG-area = 0 μm^2^, while only 15% have SG-ares > 0 μm^2^. Median values are shown as colored data points. (D) Experimental variation of median SG counts per cell and areas of single SGs that were calculated for all GFP^+^-cells between the 25% and 75% quantiles of each cell type per experiment.

**Figure S10:**
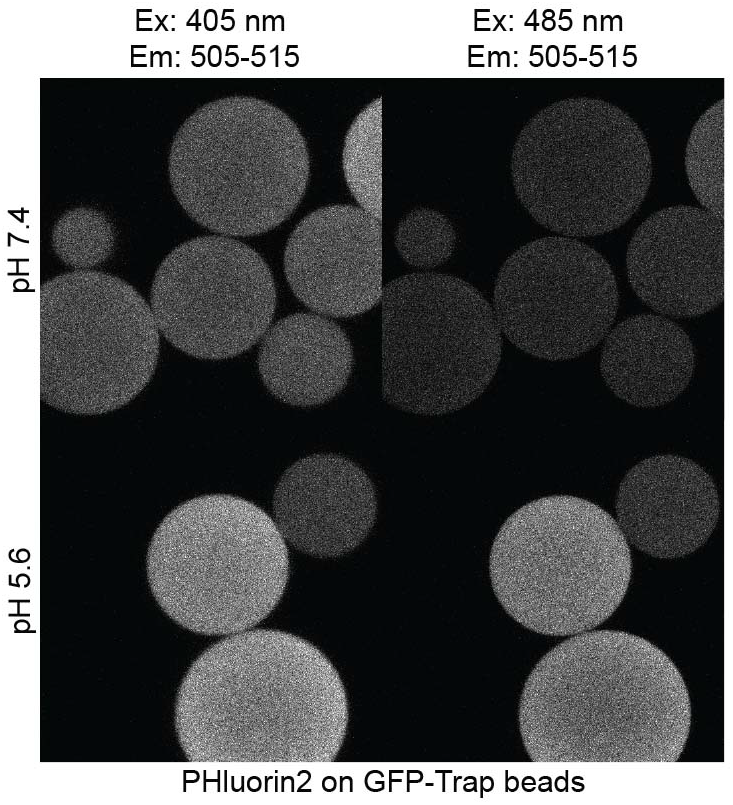
pHluorin2 bound to beads to record standard curve. Representative images of GFP-Trap agarose beads with bound GFP-pHluorin2 in pH specific buffer 7.4 (top row) and 5.6 (bottom row). Beads were sequentially excited at two excitation wavelengths of 405 and 485 nm, and the emission was detected between 505-515 nm.

### Detailed interaction description (Figures 3 and S5)

#### Site 1

In more detail, both Caprin-1^356-386^-Phe-372 and nsP3-Phe-451 are embedded in a mostly hydrophobic and aromatic environment created by NTF2 residues Val-11, Phe-15, Phe-33, Leu-114 (Cap only), Phe-124, as well as Gln-18 and the aliphatic CH2-groups of Glu-14. Residues Caprin-1^356-386^-Tyr-370 and nsP3-Leu-449 are coordinated by NTF2-Leu-10 (Cap only), Val-11 and Pro-6 as well as Asn-122 and the aliphatic CH2-groups of Glu-14 aliphatic CH2-groups of Glu-14. Residues Caprin-1^356-386^-Asn-371 and nsP3-Thr-450 are in contact with Lys-123, Asn-122 and Phe-124 (Cap only). Furthermore, the backbone amides of Caprin-1^356-386^-Asn-371 are fixed through hydrogen bonds with backbone amides of Lys-123 and Phe-124. Residues Caprin-1^356-386^-Ile-374 and nsP3-Phe-454 are in contact with Leu-22 (nsP3 only), Arg-32, Phe-33, and Lys-123. Furthermore, the backbone carbonyl of Caprin-1^356-386^-Ile-374 forms an h-bond with the amino group of Lys-123. In our pull-down assays Alanine substitutions of Caprin-1-Phe-372 and −Ile-374 abolished binding of Caprin-1 to G3BP1 completely (Figure 2B). While Ala-substitution of Caprin-1-Tyr-370 significantly reduced binding, substitution of Asn-371 did not reduce binding, suggesting Alanine as tolerated residue at this position.

#### Site 2

Although Caprin-1^356-386^ and nsP3^449-473^ cover the same secondary binding site along the NTF2 aI helix, their N- and C-terminal parts are aligned in opposite directions, respectively. Caprin-1^356-386^- Gln-365 and -Met-366 are coordinated by NTF2-residues Arg-17, Gln-18, Thr-21, Leu-22 and Gln-25. Their interactions are further strengthened by two hydrogen bonds between the side chain amides of Gln-365 and Gln-18, as well as between the backbone carbonyl of Gln-365 and the guanidine-group of Arg-17. A stretch of six residues from Glu-458 to Leu-462 of nsP3^449-473^ is coordinated by the same NTF2-residues as observed for Caprin-1^356-386^ and additionally comprises Met-29 and Arg-32. Furthermore, the carboxylate side-chain group of Asp-460 forms an ionic interaction with the guanidine-group of Arg-17. Interestingly, structural modeling puts Phe-18 of USP10 in the same position as Val-459 of nsP3^449-473^, suggesting that also USP10 might cover the second binding site along helix **a**1. Surprisingly, Ala-substitution of Gln-365 and -Met-366 did not reduce binding in pull-down assays, while Ala-substitutions of the preceding Caprin-1^356-386^-Leu-362 and Met-363 abolished binding completely. However, Leu-362 and Met-363 were not resolved in the crystal structure, thus preventing us from drawing any structural conclusions about this observation.

### Protein constructs for crystallization and biophysical measurements

>His6-TEV-G3BP 1 *-**NTF2***

MHHHHHHSSGVDLGT*<u>ENEEFQ</u>*-

***SMVMEKPSPLLVGREFVRQYYTLLNQAPDMLHRFYGKNSSYVHGGLDSNGKPADAVYGQKEIHRKVM SQNFTNCHTKIRHVDAHATLNDGVVVQVMGLLSNNNQALRRFMQTFVLAPEGSVANKFYVHNDIFRY QDEVFG***

>twSTII-<u>TEV</u>-GFP-***USP10_1-28_** // -^F3AF6A^**USP10_1-28_** //* **-Cap^356-386^** // ^F3AL9A^***Cap*_356-386_**

MSAWSHPQFEKGGGSGGGSGSAWSHPQFEKSGG*<u>ENLYFFQ</u>-*

<u>G</u>GGSVSKGEELFTGVVPILVELDGDVNGHKFSVSGEGEGDATYGKLTLKFICTTGKLPVPWPTLVTTLT YGVQCFSRYPDHMKQHDFFKSAMPEGYVQERTIFFKDDGNYKTRAEVKFEGDTLVNRIELKGIDFKED GNILGHKLEYNYNSHNVYIMADKQKNGIKVNFKIRHNIEDGSVQLADHYQQNTPIGDGPVLLPDNHYL STQSALSKDPNEKRDHMVLLEFVTAAGITLGMDELYKSENQG-

…MALHSP<u>QYIFGDF</u>SPDEFNQFFVTPRSS

*…MALHSPQYI<u>A</u>GD<u>A</u>SPDEFNQFFVTPRSS*

*… RQRVQDLMAQMQGP<u>YNFIQDSML</u>DFENQTLD*

*… RQRVQDLMAQMQGPYN<u>AI</u>QDSM<u>A</u>DFENQTLD*

### Supplemental Tables

**Table S1.** Crystallographic data table

| PDB | 6TA7 |
| --- | --- |
| Resolution range | 46.5 - 1.93 (2.00 - 1.93) |
| Space group | P 21 21 21 |
| Unit cell | 81.6 100 113.3 / 90 90 90 |
| Total reflections | 898968 (58966) |
| Unique reflections | 66520 (4630) |
| Multiplicity | 13.5 (12.7) |
| Completeness (%) | 94 (66) |
| Mean I/sigma(I) | 14.6 (0.7) |
| Wilson B-factor | 40 |
| R-meas | 0.12 (3.4) |
| CC1/2 | 1 (0.33) |
| CC* | 1 (0.70) |
| Reflections used in refinement | 66502 (4630) |
| Reflections used for R-free | 2098 (146) |
| R-work | 0.21 (0.38) |
| R-free | 0.25 (0.43) |
| CC(work) | 0.96 (0.61) |
| CC(free) | 0.92 (0.51) |
| Number of non-hydrogen atoms | 6524 |
| number_macromol | 6246 |
| number_ligands | 3 |
| Protein residues | 792 |
| RMS(bonds) | 0.007 |
| RMS(angles) | 0.78 |
| Ramachandran allowed (%) | 99.8 |
| Ramachandran favored (%) | 97 |
| Rotamer outliers (%) | 1.2 |
| Clashscore | 5.28 |
| Average B-factor (ADP) | 61 |
| ADP macromolecules | 61 |
| ADP ligands | 54 |
| ADP solvent | 55 |

**Table S2.**
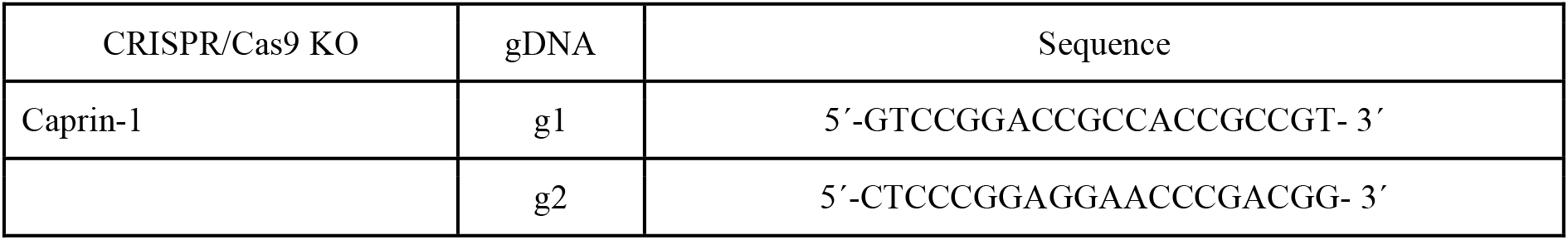
CRISPR/Cas9 sequences

**Table S3.**
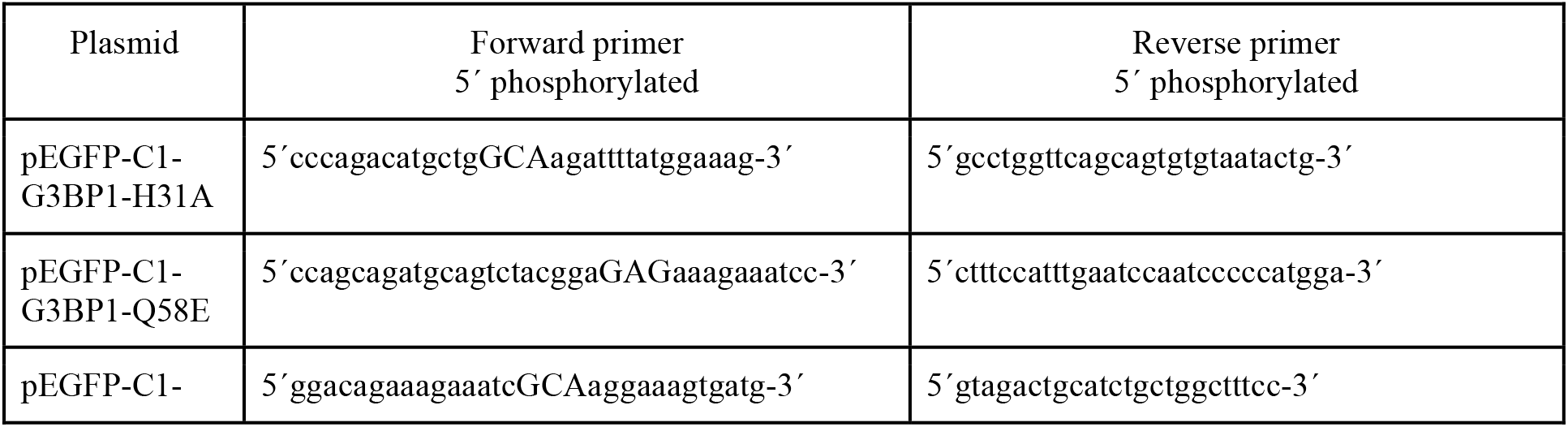
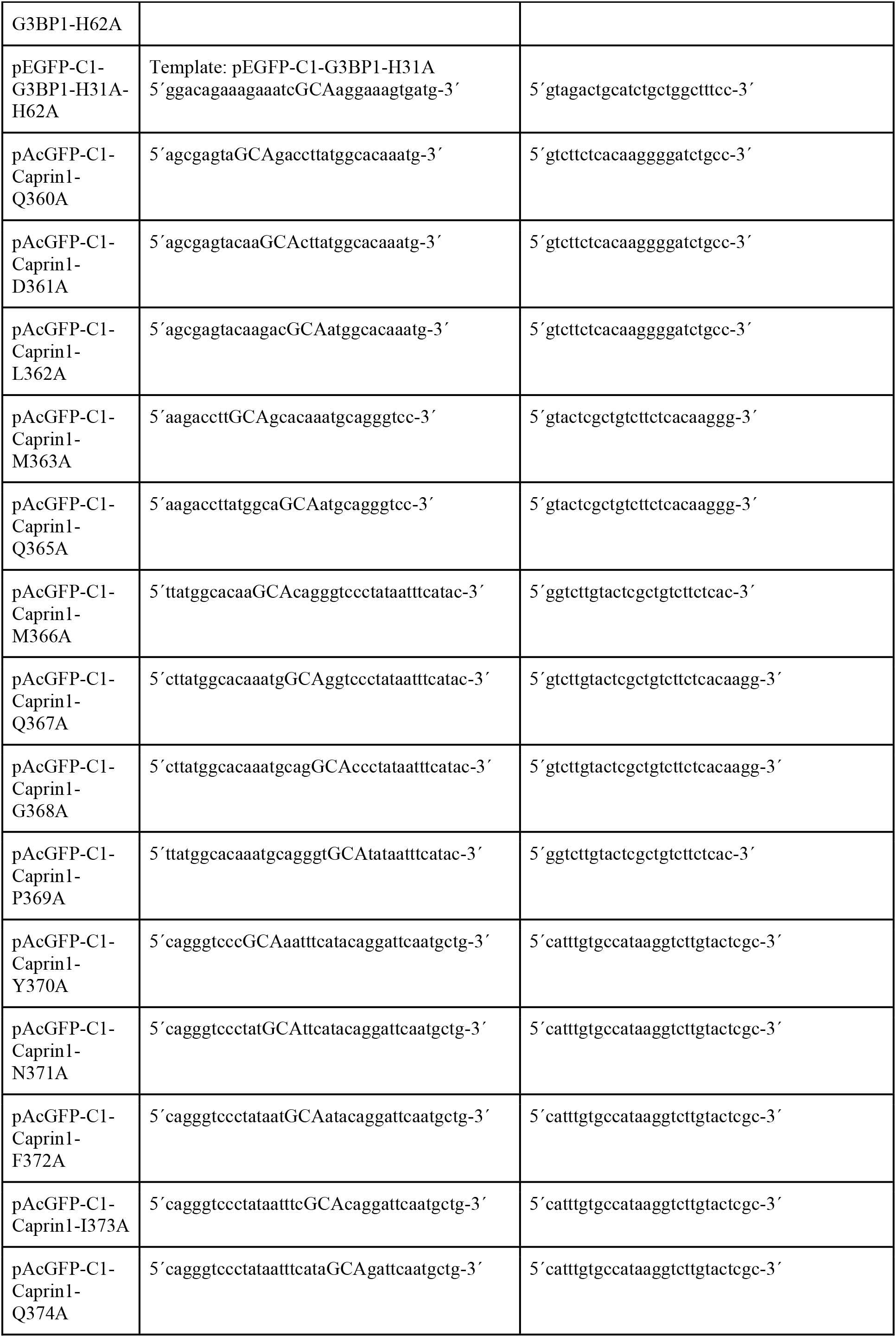
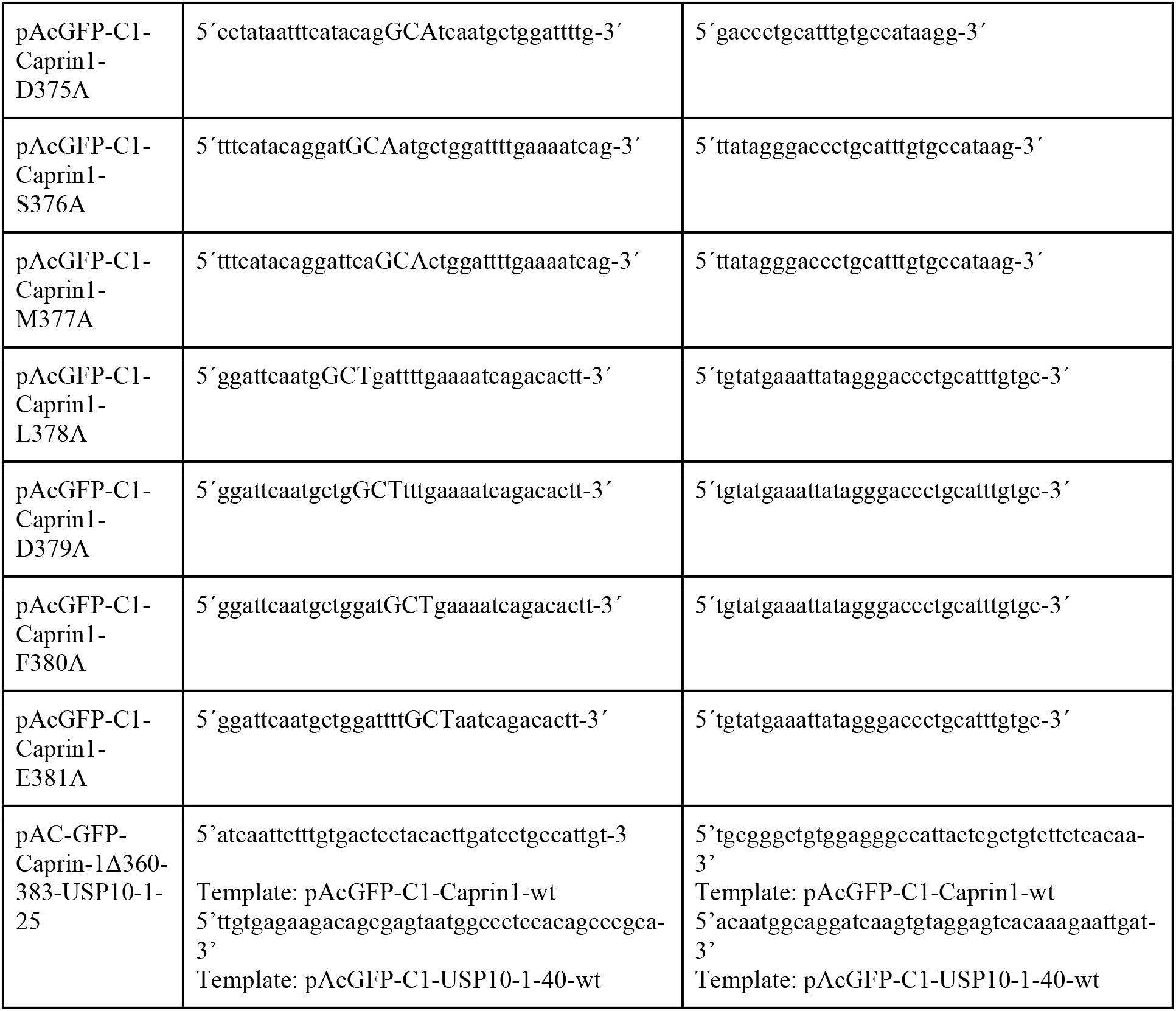
primer list

**Table S4.** Antibodies used in this study

| Antigen | Species | Catalog number | Source |
| --- | --- | --- | --- |
| Actin | Mouse | Sc-8432 | Santa Cruz Biotechnology, Inc. |
| Caprin-1 | Rabbit | 15112-1-AP | Protein Tech Group |
| Caprin-1 | Rabbit | HPA018126 | Sigma-Aldrich |
| FMR1 | Rabbit | 13755-1-AP | Protein Tech Group |
| GAPDH | Mouse | Sc-47724 | Santa Cruz Biotechnology, Inc. |
| G3BP1 | Mouse | Sc-365338 | Santa Cruz Biotechnology, Inc. |
| G3BP2 | Rabbit | C18193 | Assay Biotechnology |
| TIA-1 | Goat | Sc-1751 | Santa Cruz Biotechnology, Inc. |
| USP10 | Rabbit | A300-900A | Bethyl Laboratories |

## Footnotes

1 Karplus, P Andrew, and Kay Diederichs. 2012. “Linking Crystallographic Model and Data Quality.” *Science* Karplus, P. Andrew, and Kay Diederichs. 2015. “Assessing and Maximizing Data Quality in Macromolecular Crystallography.” *Current Opinion in Structural Biology* 34 (October): 60–68. https://doi.org/10.1016/j.sbi.2015.07.003. Adams, Paul D, Ralf W Grosse-Kunstleve, Li Wei Hung, Thomas R Ioerger, Airlie J McCoy, Nigel W Moriarty, Randy J Read, James C Sacchettini, Nicholas K Sauter, and Thomas C Terwilliger. 2002. “PHENIX: Building New Software for Automated Crystallographic Structure Determination.” *Acta Crystallographica. Section D, Biological Crystallography* 58 (Pt 11): 1948–54. Kabsch, W. 1993. “Automatic Processing of Rotation Diffraction Data from Crystals of Initially Unknown Symmetry and Cell Constants.” *Journal of Applied Crystallography* 26 (6): 795—800. https://doi.org/10.1107/S0021889893005588. Michael Krug, Manfred S. Weiss. 2012. “XDSAPP: A Graphical User Interface for the Convenient Processing of Diffraction Data Using XDS.” *Journal of Applied Crystallography* 45 (3). https://doi.org/10.1107/S0021889812011715.

2 Kelley, Lawrence A, and Michael J E Sternberg. 2009. “Protein Structure Prediction on the Web: A Case Study Using the Phyre Server.” *Nat. Protocols* 4 (3): 363–71. https://doi.org/10.1038/nprot.2009.2. London, Nir, Barak Raveh, Eyal Cohen, Guy Fathi, and Ora Schueler-Furman. 2011. “Rosetta FlexPepDock Web Server-High Resolution Modeling of Peptide-Protein Interactions.” *Nucleic Acids Research* 39 (July): W249–53. https://doi.org/10.1093/nar/gkr431. Gouet, P, E Courcelle, D I Stuart, and F Métoz. 1999. “ESPript: Analysis of Multiple Sequence Alignments in PostScript.” *Bioinformatics (Oxford, England)* 15 (4): 305–8.

